# HiSpoT: a novel one-tube dual-barcode PCR method for NGS spoligotyping amplicon library preparation in *Mycobacterium tuberculosis* complex

**DOI:** 10.64898/2026.11.04.764218

**Authors:** Paul J. Freidlin, Mor Rubinstein, Ephraim Fass, Inna Kutikov, Gal Zizelski Valenci, Yelena Losev, Zeev Dveyrin, Efrat Rorman, Israel Nissan

**Author notes:** Corresponding author: Israel Nissan, PhD.

## Abstract

Tuberculosis remains one of the leading infectious causes of death globally, making rapid, accurate, and affordable strain typing essential for effective outbreak tracking and epidemiological surveillance. To address the limitations and high costs of current typing methods, we developed a novel one-tube, dual-barcode PCR method for preparing targeted next-generation sequencing (NGS) amplicon libraries for spoligotyping of Mycobacterium tuberculosis complex (MTBC), which we termed HiSpoT (High-throughput Spoligotyping by Targeted next generation sequencing). The assay targets 43 conserved spacer oligonucleotides in the CRISPR direct repeat (DR) region and is presented here together with the previously unpublished, difficult-to-design dual-barcode primer set needed to multiplex 96 samples in a single tube. HiSpoT is accurate, fast, high-throughput, and low-cost, eliminating the need for a commercial NGS library preparation kit. Because HiSpoT is PCR-based, the resulting spoligotypes remain directly comparable with the large PCR-based spoligotype databases accumulated over decades, enabling rapid outbreak tracking and biogeographic characterization of a country’s tuberculosis burden while higher-resolution whole-genome sequencing (WGS) data are obtained. Analysis of 288 clinical MTBC isolates from Israel (2020-2022) demonstrated a broad range of genetic lineages, and comparison of 79 isolates analyzed by both HiSpoT and WGS showed 70% full concordance, with essentially all discrepancies attributable to single-spacer insertion-element disruptions of PCR primer sites. We also present two open-source Google Colab Python tools, Spoligotyping_MTBC_gcl and its condensed HiSpoT_Spoligotyping_v8_ready VIBE variant, freely available on GitHub, which automatically process Illumina paired-end fastq output from HiSpoT or WGS and report binary and octal spoligotypes, spoligo international types (SIT), and MTBC family assignments with accompanying quality-control reports.

**IMPORTANCE:** Tuberculosis, caused by *Mycobacterium tuberculosis*, is the leading cause of death from a single infectious agent worldwide, and rapid, accurate, and affordable strain typing is essential for tracking outbreaks and directing limited public health resources. The decades-old laboratory method for spoligotyping, a widely used genetic fingerprint for this bacterium, is slow, labor-intensive, and hard to read. Newer sequencing-based approaches solve these problems but usually still need an expensive, time-consuming commercial kit to prepare samples for sequencing. This work describes HiSpoT, a simplified, low-cost, one-tube method that prepares samples for sequencing without a commercial kit, cutting turnaround from about a week to about two days while testing many samples together. Because HiSpoT results agree with results obtained from older typing methods, laboratories anywhere with basic sequencing equipment can compare results to decades of stored international data. Two free, open-source computer programs are also provided so laboratories with limited bioinformatics expertise can analyze their sequencing data automatically.

## INTRODUCTION

### Importance of HiSpoT: **Hi**gh-throughput **Spo**ligotyping by **T**argeted next generation sequencing

According to a recent WHO report (1), tuberculosis is the number one cause of death by a single infectious agent, and one of the top ten causes of death globally. In 2024 more than a million people died of tuberculosis. Tuberculosis is found in all countries (2), in people of all ages and although treatable, tuberculosis poses a major public health challenge due to the development of multiple drug resistance (3), relatively long treatment times, and worldwide spread due to the rise of global tourism and large-scale immigration from endemic-risk countries (4, 5). Rapid, sensitive, accurate, affordable high-throughput targeted (aka focused) genotyping obtainable with HiSpoT is a priority in order to quickly track and limit tuberculosis outbreaks in developed and resource-poor countries, and simultaneously characterize a country’s biogeographic tuberculosis burden in order to prioritize the utilization of limited public health resources. The fact that HiSpoT is a PCR-based assay, makes HiSpoT spoligotypes compatible with major international databases historically constructed with thousands of PCR-based spoligotypes (6, 7, 8).

HiSpoT is a fast, sensitive, accurate, high-throughput, low-cost, labor-saving method for genotyping tuberculosis by targeted NGS spoligotyping of *Mycobacterium tuberculosis* bacterial complex (MTBC). HiSpoT features a novel platform for one tube dual-barcode PCR for preparation of a ready to use targeted NGS amplicon library. An important benefit is that the costly commercial kit for preparation of NGS libraries is not required, which in addition to materials’ cost savings, eliminates a labor and time intensive step in the sequencing protocol. We announce the HiSpoT method in this paper, and through an open-access royalty-free patent at https://patents.google.com/patent/US20210355526A1/en. In addition, we present the software we developed to analyze the MiSeq Illumina HiSpoT targeted NGS output. Our software also can analyze MTBC WGS NGS output. We make our software available through downloading from GitHub.

### Spoligotyping

Soon after the discovery of CRISPR in *E. coli* (9), CRISPR in MTBC was discovered and quickly exploited for molecular epidemiology. A robust PCR-based reverse line blot was developed and commercialized, and the 43 spacer spoligotype became the standard for MTBC genotyping (10, 11, 12, 13) until additional genotyping techniques were available for more precise molecular epidemiology (7, 14, 15, 16). The reverse line blot technique is labor intensive, and the results, spots on film produced by chemiluminescence, are sometimes difficult to interpret, and manually time-consuming to score and report, creating opportunities for typos (incorrectly written results). Other non-NGS techniques for spoligotyping have been developed (17, 18, 19, 20, 21), but require special equipment and skills, as opposed to the now widespread availability of NGS platforms and trained technical staff.

The presence or absence of each of 43 highly conserved MTBC-associated unique spacer oligonucleotides is a deep phylogenetic trait of MTBC families, with significant correspondence to the large sequence polymorphisms (LSPs) characteristic of MTBC families (14, 22, 23, 24, 25, 26) and the SNP-derived MTBC lineages (14). The MTBC CRISPR/DR locus consists of 36-bp repeated sequences (direct repeats, DRs) that are interspersed by 34-41-bp non-repetitive spacers (10, 11, 12, 13). The spacers are derived from phage or plasmid sequences that have been integrated into the MTBC genome as a result of ancient infection events resolved via CRISPR-CAS adaptive acquired bacterial immunity (27, 28, 29, 30). The number and order of spacers vary among MTBC strains, reflecting their evolutionary history and genetic diversity (12, 22, 31).

The spacers are flanked by CRISPR repeat sequences highly conserved among the MTBC, thus PCR amplification of the 43 chosen MTBC spacers for all CRISPR-containing MTBC has been accomplished by the use of repeat-based sequences as primers (11). The result, the spoligotype, is given as a 43 consecutive binary expression of 0’s (spacer absent) and 1’s (spacer present) summarizing the spacer status of any given MTBC clinical isolate. Alternatively, by convention the 43-digit binary expression is converted to a fifteen number octal expression of 14 octal numbers derived consecutively from groups of three binary expression numbers, with the last number 15 being 0 (absent) or 1 (present) (32) (Fig. 1).

**FIG 1.**
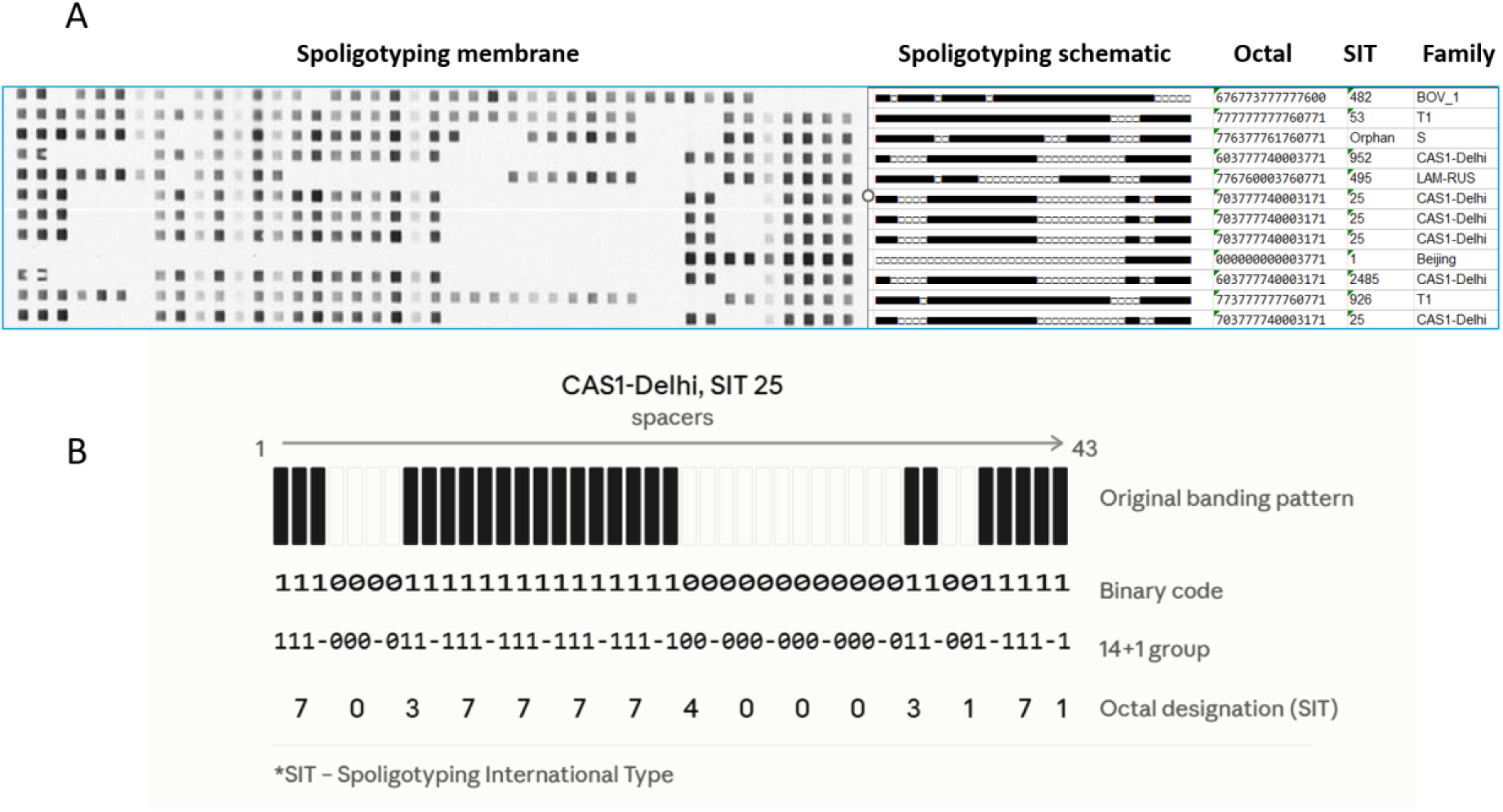
A. Conventional membrane-based spoligotyping of *Mycobacterium tuberculosis* complex (MTBC). A typical spoligotyping hybridization membrane showing the 43 spacer positions (numbered 1 to 43); black spots indicate hybridization (spacer present, binary “1”), and blank positions indicate absence (binary “0”). B. An example binary spoligotype of CAS1-Delhi family, SIT 25 is converted, in successive groups of three digits (14 groups plus a final single digit), into the 15-digit octal spoligotype international type (SIT) code. The conventional workflow used to generate this result is shown in Fig. 3.

Spoligotyping serves as a gold standard for MTBC molecular epidemiology and separation of biogeographic lineages. A spoligotype dissimilar from a prototypic outbreak spoligotype conclusively rules out membership in that outbreak, while a spoligotype identical to the outbreak spoligotype flags a need for higher resolution molecular epidemiologic analysis. Thus, HiSpoT quickly identifies putative outbreak members which can then be given priority in the whole genome sequencing (WGS) workflow. Large international databases existed for PCR-based spoligotypes (2, 6, 8) and our software accesses a version published in 2016. Currently (at publication date of this article) at least two spoligotype databases are available:

https://www.mbovis.org/database.php

https://tbinsight.cs.rpi.edu/run_tb_lineage.html

Since HiSpoT is PCR-based, the resultant spoligotypes obtained by HiSpoT can reference the international databases (such as our 2016 version) to help characterize the evolutionary (phylogenetic) and biogeographic sources of a country’s tuberculosis burden. Many countries, including resource-challenged nations, have created NGS centers which task whole genome sequencing (WGS) of microorganisms including MTBC. WGS of MTBC is slower and more costly than HiSpoT, but importantly provides single nucleotide polymorphism (SNP)-based clonal identity of MTBC for conclusive molecular epidemiology of outbreak tracing, and provides drug resistance genotypes for quick decisions on treatment modalities (14, 16, 33, 34, 35).

Spoligotypes can also be obtained from WGS whole genome sequences (36) or WGS paired-end read sets (37). Both Google Colab based HiSpoT_Spoligotyping_v8_ready and Spoligotyping_MTBC_gcl softwares (which we have provided for downloading from GitHub) can extract spoligotypes from WGS paired end read-sets (approximately 1 minute for HiSpoT_Spoligotyping_v8_ready, 8 to about 10 minutes for Spoligotyping_MTBC_gcl softwares for each WGS paired-end fastq read set). For the smaller HiSpoT targeted NGS files of paired end read sets analysis takes about 30 seconds for 96 paired-end read sets using HiSpoT_Spoligotyping_v8_ready or Spoligotyping_MTBC_gcl softwares. Any country with NGS facilities can perform HiSpoT which is much faster, more high-throughput, lower-cost, and labor-saving than WGS and alternative PCR-based spoligotyping techniques. In addition, spoligotypes generated by HiSpoT give quick, sometimes conclusive, molecular epidemiologic information concerning outbreak membership, clear deep phylogenetic status, and data which allow biogeographic categorization.

## MATERIALS AND METHODS

### Development and Background of HiSpoT

HiSpoT evolved from the development of various approaches to validate the results of the rare complete absence of MTBC DR region spacers (deleted DR) in reverse line blot spoligotyping (22). In its initial form, the assay known as “HiSpo targeted NGS” used a commercial library preparation kit. Empirical and theoretical work allowed the design and validation of probes, primers, and barcodes that avoided intra and inter non-specific hybridization events, and that performed at the desired melting temperatures. HiSpo Illumina MiSeq output was analyzed by an early stand-alone python version of the current Spoligotyping_MTBC_gcl software. Subsequent evolution of the software was based on 100% agreement of the new version with the previous version’s results. Approximately 97 percent of the stand-alone HiSpo spoligotype results agreed with the reverse line blot results (data not shown). Cases of poor signal weak reverse line blot chemiluminescence were resolved by clear good signal HiSpo results for the spacer in question. Subsequent work built on the HiSpo platform to create the HiSpoT one tube, dual-barcode method that eliminated the need for the commercial library preparation kit.

### HiSpoT Materials, Protocol References, and Safety

Primers, barcodes, and assay preparation and performance instructions for up to 96 samples and negative and positive controls, are found in supplementary information SI_1, SI_2, and SI_3. If you need help in setting up HiSpoT in your facility, please contact Dr. Israel Nissan at who will try to help you overcome whatever difficulties you have encountered. All work with MTBC should be done in a BSA 3 hood until after heat inactivation. For HiSpoT and WGS, DNA was prepared robotically using Promega Maxwell 16 following manufacturer’s instructions.

### Bioinformatic Pipeline Logic and Threshold Determination Software Analysis Pipeline and Threshold Rules

Analyses were performed with the Spoligotyping_MTBC software, modified for cloud execution as a Spoligotyping_MTBC_gcl Google Collaboratory Notebook (Jupyter, ipynb). An important advantage to the use of Spoligotyping_MTBC_gcl and the VIBE version HiSpoT_Spoligotyping_v8_ready is their ability to automatically convert “.gz” HiSpoT Illumina MiSeq output into demultiplexed paired-end fastq read sets, and automatically create and import files necessary for the analysis of the fastq files. Results are reported as a clinical report in Excel format, and as quality control reports in text format.

Software for analysis of HiSpoT and WGS paired-end fastq read set outputs, which we provide for downloading from GitHub, automatically processes the Illumina “.gz” output, and with a few steps interactive with the user, analyzes the resulting fastq paired-end read sets, and reports the results in Excel format as binary and octal spoligotype, SIT, and MTBC family, along with quality control reports in text formatted for clinical laboratory use. The python software uses partial spacer 17 mer (17 nucleotide sequences) *in silico* probes, no mismatching allowed, to search for and score the spacers in each sample. The threshold is to always consider “less than 10 matches” as negative for presence of a spacer (0), otherwise for HiSpoT targeted (aka focused) NGS: 1) less than 0.15 of the mean number of matches for the 43 spacers is considered negative (0), while greater than or equal to 0.15 of the mean is considered positive (1), and for WGS NGS: less than 0.365 of the mean number of matches for the 43 spacers is considered negative (0), while greater than or equal to 0.365 of the mean is considered positive (1). These thresholds were arrived at by some theoretical statistical considerations, but also with significant empirical input. The current VIBE version of the software (v8) utilizes a different algorithm for threshold calculation for WGS, but gives results 100% in agreement with the object oriented Python version Spoligotyping_MTBC_gcl. The user of the software to analyze HiSpoT or WGS NGS paired-end read sets for accurate spoligotypes, must first see that the software works properly with the validation read sets provided with the software. Then, with known positive and negative control user-extracted DNA, the user should validate the software results. Quality and amount of DNA could affect the results along with other variables, therefore the user must validate the software results with the user’s own gold-standard positive and negative controls before relying on the software for routine spoligotyping of the user’s own generated HiSpoT or WGS NGS paired end read sets. Another possible problem is that the software uses the Google-provided latest version of python (in this article version 3.13). Should a future python version not be back-compatible with previous versions, either access to the correct previous version would have to be done by (AI assisted) coding, or AI assisted modified coding of the main program would have to be done so it would work with the new not back-compatible version of python.

### Software installation Spoligotyping_MTBC_gcl

Spoligotyping_MTBC_gcl software is available to download from GitHub as a python ipynb (interactive python notebook) Jupyter notebook file, which when uploaded to your Google Drive, is used as a Google Colab Notebook. First, create a folder (aka directory) in your Google Drive which you name “data,” and another folder which you name “Colab Notebooks.”

Next, download to your computer from GitHub the Spoligotyping_MTBC_gcl software. The Spoligotyping_MTBC_gcl_ver01.ipynb software should then be uploaded to your Google Drive folder “Colab Notebooks,” and 3 folders each of which should be uploaded to the “data” folder in your Google Drive. The folder “spo_software_modules_and_template_file” contains two python modules and an Excel template which are utilized by the spoligotyping_MTBC_gcl_ver01 Colaboratory notebook to analyze and summarize HiSpoT, or WGS, Illumina paired end read sets in an object-oriented manner. The notebook and accompanying files are copyright 2025 by the Government of the State of Israel under the MIT license. The remaining 2 folders contain validation data which you will access and analyze before using the software to analyze your own data. The folder “HiSpoT_validation_data” contains 5 Illumina paired-end fastq read sets output. If the files are in compressed .gz format, the program will still work. The HiSpoT results for validation comparison are given in the folder “HiSpoT_results_spoMTBC_on_five_strains.” We suggest analyzing the WGS paired end read set T3_0525-11 which we previously submitted to GenBank (22).

### HiSpoT_Spoligotyping_v8_ready_kit

This is an AI-VIBE compressed version of the original Spoligotyping_MTBC software.

If not already present, create a folder “data” in your Google Drive, and folder “Colab Notebooks.”

Download from GitHub the HiSpoT_Spoligotyping_v8_ready.ipynb software into your folder “Colab Notebooks.”

Download from GitHub the user manual and validation data and results. Follow the detailed information in the user manual for setting up the program and analyzing HiSpoT fastq files or WGS fastq files (note: the program automatically detects whether the paired end read set is HiSpoT or WGS).

## RESULTS

### Development and Workflow of the Novel HiSpoT Method

To create a streamlined, cost-effective alternative to commercial library preparation kits, we designed a novel single-tube dual-barcode PCR strategy. The basic set-up of HiSpoT is shown in Figure 2. In this step-wise, temperature-dependent protocol, sample specific DNA or negative or positive test controls are added separately to each of 96 microplate wells containing the appropriate Hotstart PCR enzyme - primers mix. First, to each well is added a mix containing primers specific for the DR repeats in the MTBC CRISPR/DR region, the primers being connected to universal tails (UT). Then to each separate well (separate sample or negative or positive control wells) – are added primers consisting of UT specific complementary sequences attached to sample specific dual barcode sequences (for sample labeling) and further attached to Illumina adapter sequences (P5, P7) for flow cell binding.

**FIG 2.**
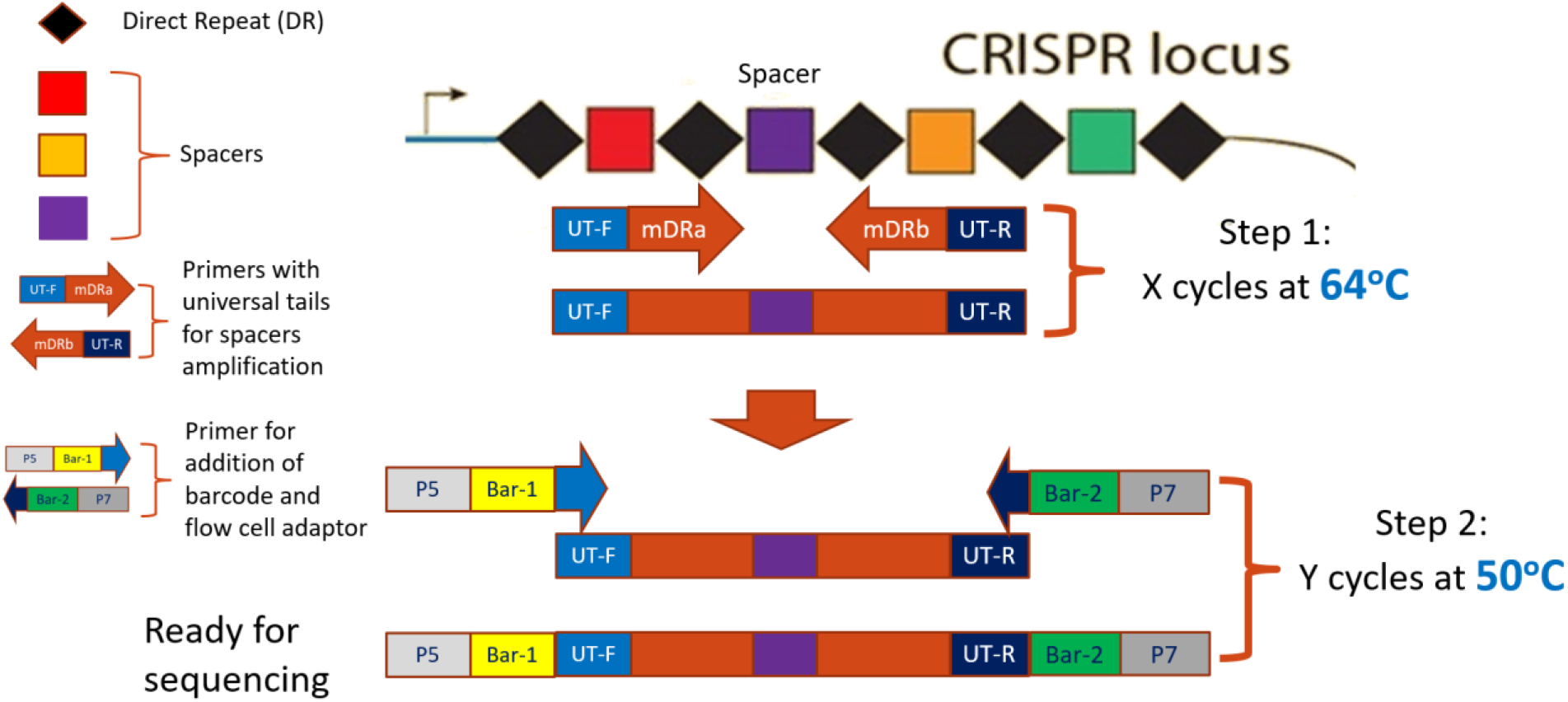
Principle of operation of the HiSpoT single-tube dual PCR (std-PCR) method. All primers required for library preparation are pre-loaded into a single reaction tube; selective primer activation is achieved by annealing temperature alone. Step 1 (X cycles at 64°C): primers bearing universal tails (UT-F/UT-R) and MTBC direct-repeat-region-specific sequences (mDRa/mDRb) amplify the CRISPR spacers exclusively. Step 2 (Y cycles at 50°C): a second primer set anneals to the universal tails introduced in step 1, adding sample-specific dual barcodes (Bar-1, Bar-2) and Illumina flow-cell adapters (P5, P7). The resulting amplicons are pooled, cleaned, and are directly ready for sequencing, without a separate commercial library preparation step or normalization.

Restoring base diversity in a low-complexity amplicon library: every molecule in a CRISPR amplicon pool begins with the same conserved direct repeat, so the first sequencing cycles are near-monomorphic - a problem for Illumina chemistry, which depends on base diversity for cluster calling and phasing. To solve this problem, our inner primers carry a variable-length degenerate insert of zero to three N bases between the universal tail and locus-specific sequence; so each cycle generates a mixed base composition - restoring complexity without changing the amplicon, workflow or analysis, reducing the amount of PhiX and increasing target yield.

The first PCR reactions are run at the elevated temperature of 64 degrees C at which only the DR-UT primers can bind and initiate PCR on the MTBC bacterial chromosome. Proofreading KOD Hot Start DNA polymerase (cat No. 71086) is used to generate a high-quality double stranded blunt-end DNA library. LightCycler 480 Resolight Dye (Cat. No. 04909640001) is added to the reaction mixture to enable real time monitoring of individual library success by PCR amplification and melting curve. Cycling at 64 degrees C exhausts the supply of DR-UT primers, at which point all spacers present in the DR region are contained in amplicons attached to DR-UT primers. Subsequently, the second PCR set of cycles is performed at the lower temperature of 50 degrees C suitable for the “UT-complement - sample specific dual barcode - flow cell adapter” primers to anneal to the UT tails of the spacer containing amplicons made in the first set of cycles. This second PCR reaction at 50 C amplifies the spacer-containing amplicons in each well in a sample specific (dual barcode labelled) manner. At completion, the HiSpoT PCR products are pooled, cleaned and used as the library for Illumina MiSeq NGS sequencing. Crucially, no library normalization is needed thus saving time, work and costs. The MiSeq Illumina instrument de-multiplexes the HiSpoT NGS sequences according to the appropriate dual barcode sample identification label, and outputs them in a “.gz” file of paired-end fastq read sets.

Figure 3 shows a workflow comparison of the reverse line blot and HiSpoT targeted NGS methods. The advantages of HiSpoT in reduced cost, less time and labor consumption, and automated analysis are clearly seen.

**FIG 3.**
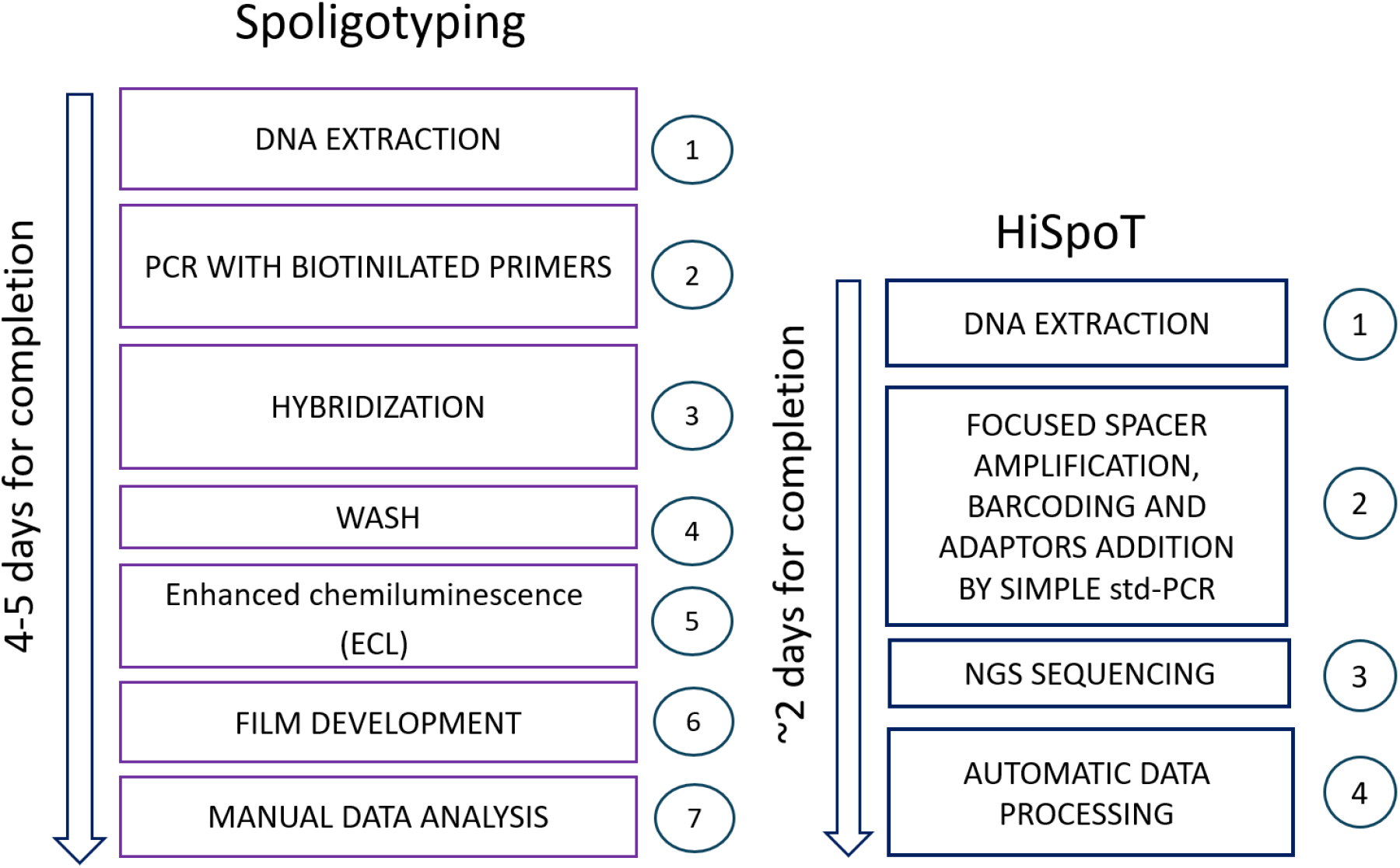
Workflow comparison between conventional membrane-based spoligotyping and the HiSpoT method. Conventional spoligotyping requires 4 to 5 days across seven manual steps (DNA extraction, PCR with biotinylated primers, hybridization, wash, enhanced chemiluminescence, film development, and manual data analysis; representative result shown in Fig. 1). HiSpoT requires approximately 2 days across four steps (DNA extraction, focused spacer amplification/barcoding/adapter addition by single-tube dual PCR, NGS sequencing, and automated data processing), enabling a larger number of samples to be processed per run with fully automated result reporting (representative output shown in Fig. 8).

### Analysis of clinical tuberculosis isolates using the HiSpoT method (Figure 4)

4A. Two hundred and eighty-eight tuberculosis samples, from the years 2020-2022, were analyzed using the HiSpoT method. A variety of genetic lineages were obtained. “Novel” indicates samples whose spoligotyping profiles were not found in the SITVIT2 international database of 2016.

**FIG 4.**
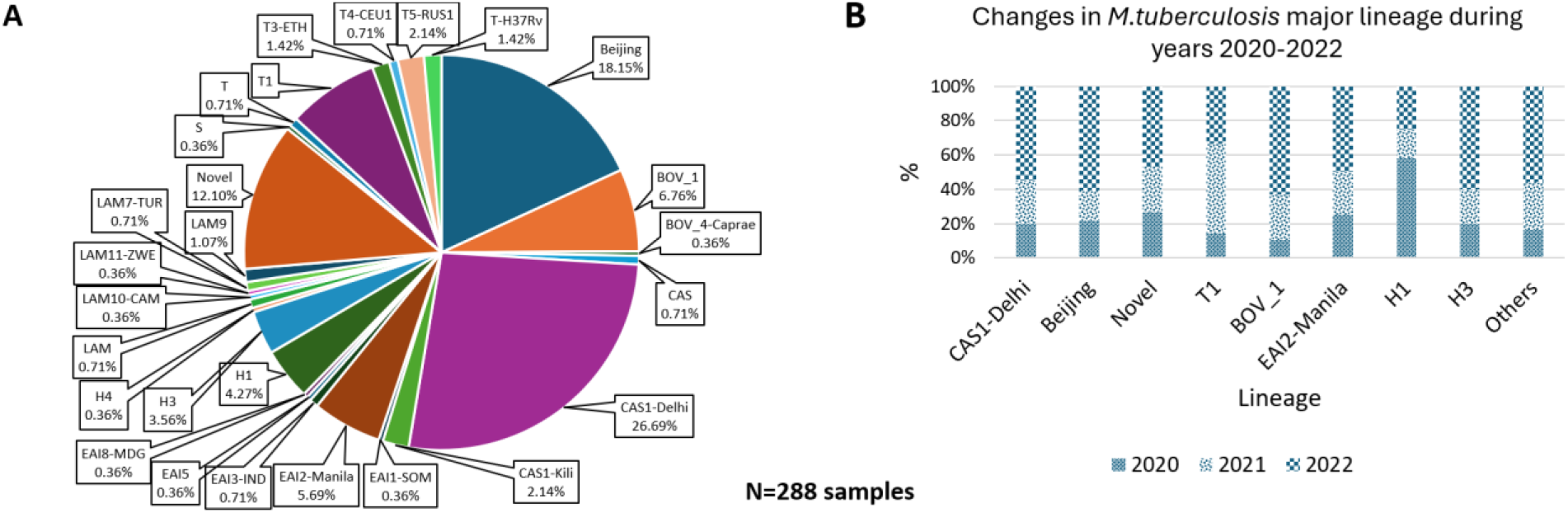
Genetic lineage distribution of *Mycobacterium tuberculosis* complex isolates in Israel, 2020-2022, determined by HiSpoT. (A) Lineage distribution among 288 clinical isolates analyzed by HiSpoT. “Novel” denotes spoligotype profiles not present in the SITVIT2 international database. (B) Year-by-year change (2020, 2021, 2022) in the relative prevalence of the eight leading lineages; “Others” comprises all remaining lineages. N = 288 samples.

4B. Figure 4B shows the change in the prevalence (in percentages) of the eight leading tuberculosis lineages in Israel in 2020-2022. “Others” indicates all other lineages.

### Real-Time PCR Monitoring via Melting Curve Analysis

To ensure library quality and filter out non-target organisms prior to sequencing, we integrated real-time melting curve monitoring into the reaction scheme.

As seen in Figure 5, melting curves give a clear indication of status of the amplification products. MTBC amplicons cluster together, whereas amplicons from negative test control water (NTC) and non-tuberculosis mycobacteria (NTM) are clearly separated from the MTBC amplicon cluster. The amplification (not presented) and melting curve data provide vital, real-time information regarding the success of each library, enabling the library preparation step to be repeated prior to NGS sequencing if necessary. This also allows MTBC and NTC amplicons to be pooled for sequencing, to the exclusion of NTMs.

**FIG 5.**
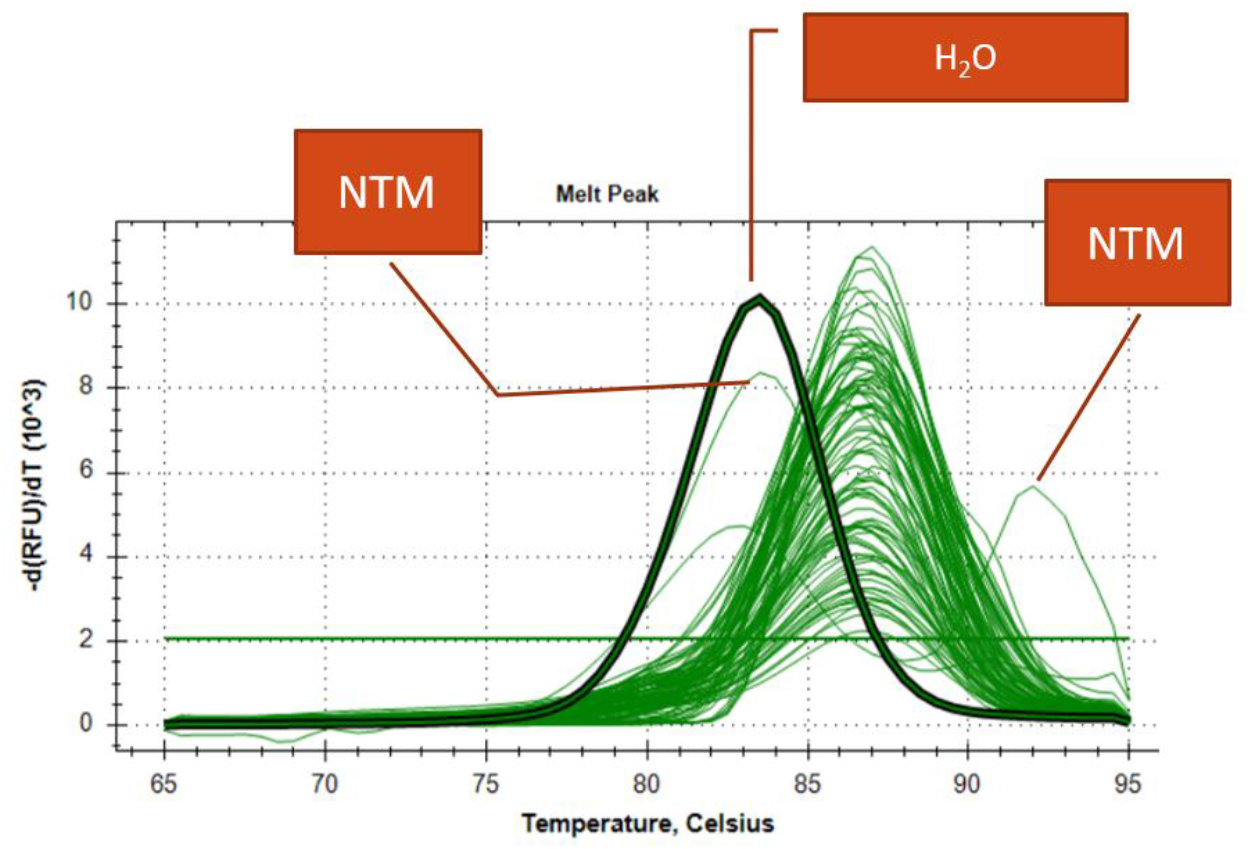
Representative HiSpoT real-time PCR melting curve analysis. MTBC-positive amplicons form a tight, characteristic melting peak cluster (bold curve highlighted), clearly distinguishable from the melting peaks of non-tuberculous mycobacteria (NTM) and the no-template (H₂O) negative control. Real-time melting curve monitoring allows library preparation failures to be identified and repeated before proceeding to NGS sequencing.

### Illumina MiSec demultiplexes HiSpoT sequencing output

Following real-time quality control, amplicons were sequenced to assess demultiplexing efficiency and signal clarity. Even if NTMs are included for sequencing, the demultiplexed results clearly differentiate among the NTMs, the NTCs, and the MTBC (Figure 6). For each of dual barcode labeled 96 samples and NTC, the graph shows a measure of the number of reads detected per dual barcode label (sample or NTC). The number of reads identified for NTC and NTM are none or close to none, thus visually clearly differentiating the NTC and NTM from the MTBC samples. It is important to note that this result is achieved without library normalization, a process that requires time and resources. This is consistent with the fact that HiSpoT provides excellent signal to noise ratio, as the example shown in Figure 7 illustrates. To obtain these results library normalization was not required, nor was a commercial kit required for library preparation. HiSpoT eliminated the expensive, time-consuming, labor intensive library preparation step, which is a strong endorsement for the use of HiSpoT for targeted NGS spoligotyping of MTBC, or for that matter, a strong endorsement for the one tube dual-barcode method as a general platform for any type of targeted NGS.

**FIG 6.**
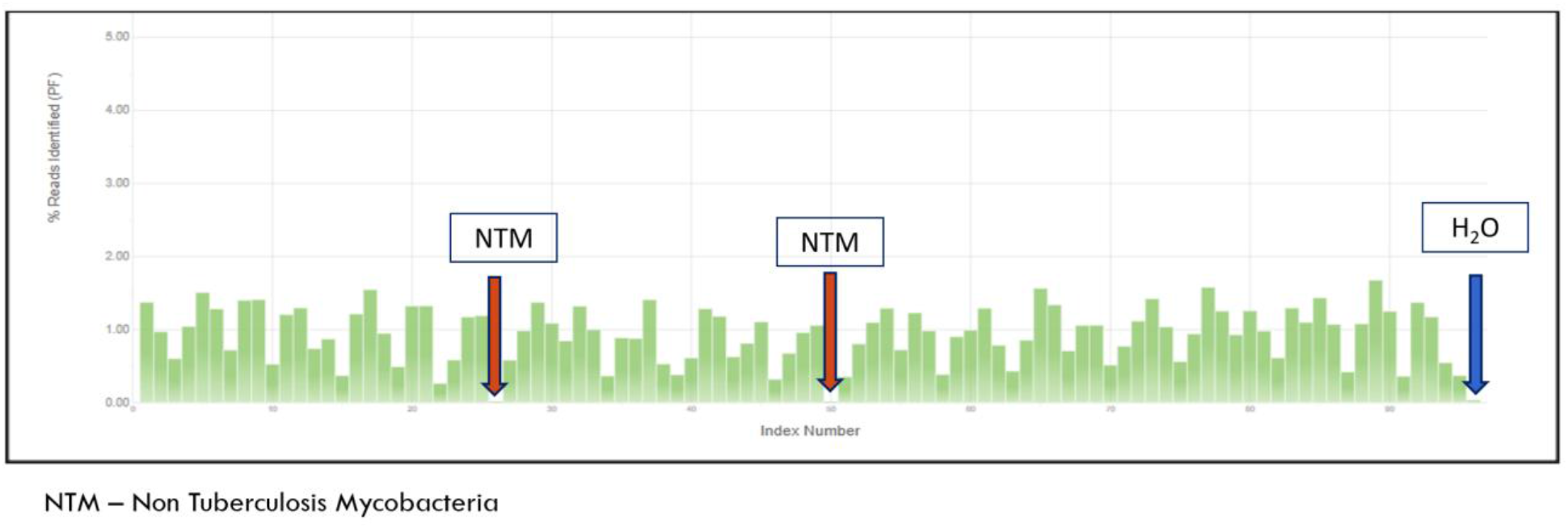
Representative Illumina MiSeq de-multiplexing output for a HiSpoT sequencing run (Nano Kit v2, approximately 16-hour run). Bars show the percentage of reads identified (passing filter) per dual-barcode index across 96 samples plus controls. Non-tuberculous mycobacteria (NTM) and the no-template water control (H₂O) show negligible read counts relative to *Mycobacterium tuberculosis* complex-positive samples, despite the absence of library normalization prior to sequencing.

**FIG 7.**
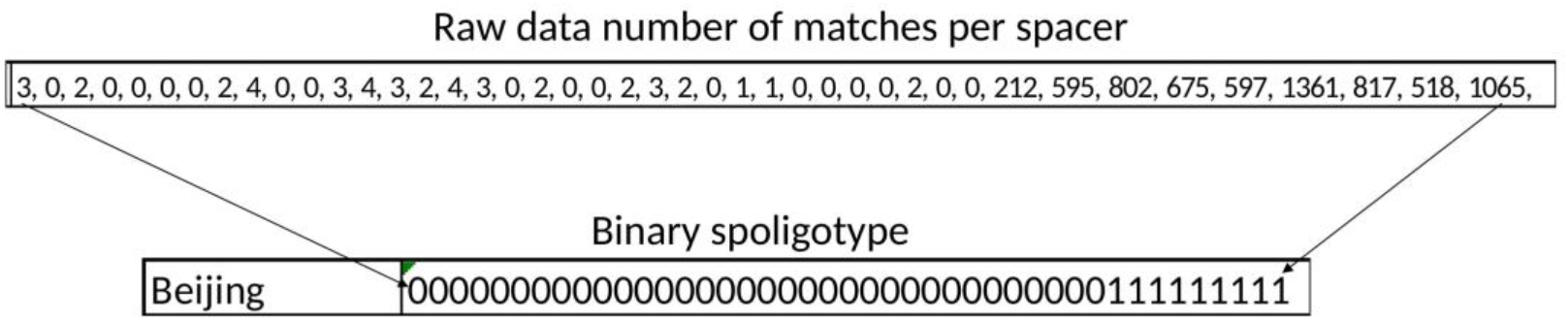
Signal-to-noise ratio of HiSpoT targeted NGS output, illustrated for a representative Beijing-lineage isolate. Top: raw number of sequence-read matches per spacer (43 values, spacers 1 to 43). Low counts correspond to absent spacers and high counts (here, several hundred to >1,000 reads) correspond to present spacers, yielding the binary spoligotype shown below.

### Reports outputted by the VIBE generated HiSpoT_Spoligotyping_v8_ready software (VIBE, that is AI processed, version of the Spoligotyping_MTBC_gcl software)

To automate read processing and clinical interpretation from demultiplexed fastq files, dedicated bioinformatics software was applied. Figure 8 presents an example of the Excel output of the VIBE generated HiSpoT_Spoligotyping_v8_ready.ipynb software. The Excel summary includes sample names, binary and octal spoligotypes, SIT and MTBC family association and in addition quality control information. This is accompanied by a text report. The Spoligotyping_MTBC_gcl.ipynb software provides similar information in Excel format with two accompanying text reports.

**FIG 8.**
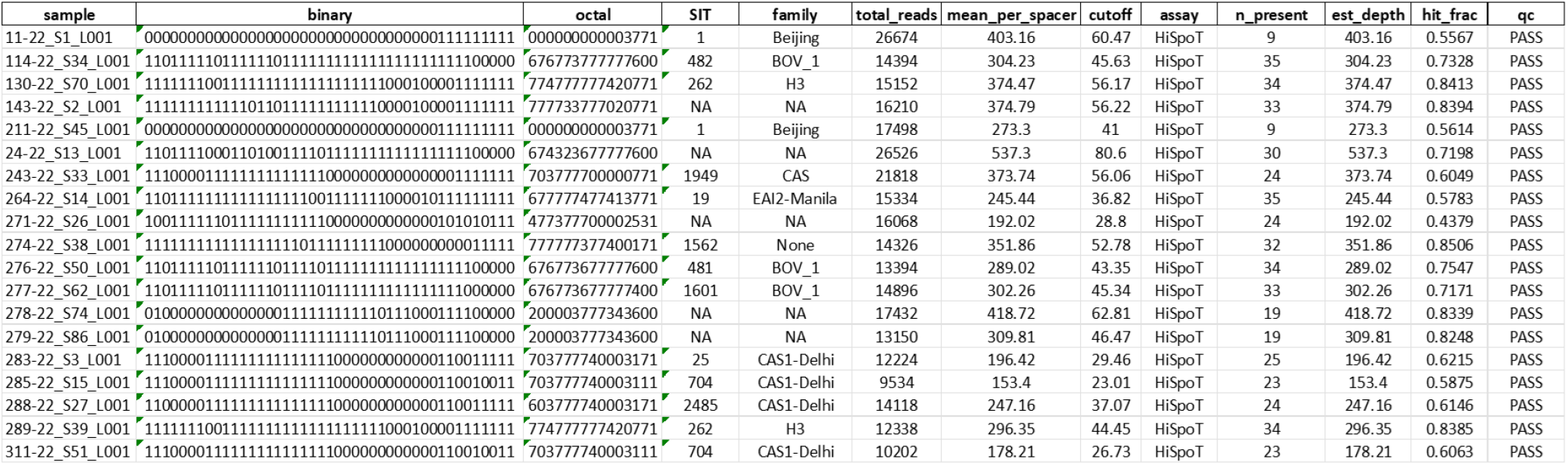
Representative output of the HiSpoT_Spoligotyping_v8_ready analysis software. Spoligotypes of 19 *Mycobacterium tuberculosis* complex isolates obtained with HiSpoT amplicon sequencing and the HiSpoT v8 analysis pipeline. Each row is one isolate, analyzed from paired-end Illumina reads (R1 + R2). sample, isolate identifier derived from the FASTQ file name. binary, presence (1) or absence (0) of the 43 standard direct-repeat (DR) spacers, listed in order from spacer 1 to spacer 43. octal, the 15-digit octal code of the spoligotype. Spacers 1–42 are converted in groups of three, and spacer 43 forms the final digit. SIT and family, the SITVIT shared international type and lineage/family assigned by exact matching of the octal code to the embedded SITVIT dictionary. NA indicates a pattern with no SIT in the dictionary. “None” indicates a SIT with no family assignment in SITVIT. total_reads, number of reads scanned (R1 + R2). mean_per_spacer, total spacer-containing read counts divided by 43. cutoff, minimum read count for a spacer to be called present, calculated as max (10, 0.15 × mean_per_spacer). assay, calling mode applied. n_present, number of spacers called present. est_depth, depth estimate used for calling; in HiSpoT mode this equals mean_per_spacer. hit_frac, fraction of reads containing at least one spacer sequence. qc, quality verdict; PASS indicates reads present, at least one spacer detected, and a mean per-spacer count of at least 15.

### Concordance between WGS and HiSpoT spoligotypes

To benchmark the accuracy of HiSpoT against whole-genome sequencing (WGS), 79 clinical isolates were analyzed in parallel by both methods. Table SI_4 compares the results from Spoligotyping_MTBC analysis of output from the PCR based HiSpoT targeted NGS of 79 clinical isolates versus the Spoligotyping_MTBC analysis of WGS NGS sequencing output from the same 79 clinical isolates.

Table 1. summarizes the results presented in Table SI_4.

**Table 1.** Summary of spacer discrepancies between whole-genome sequencing (WGS) and PCR-based HiSpoT targeted next-generation sequencing (NGS) spoligotypes for 79 *Mycobacterium tuberculosis* complex clinical isolates (2020–2021).

| Family | SIT (HiSpoT) | SIT (WGS) | No. of strains | Discrepant spacer | Spacer status: WGS | Spacer status: HiSpoT |
|---|---|---|---|---|---|---|
| CAS1-Delhi | 25 | none | 13 | 23 | present (1) | absent (0) |
| CAS1-Delhi | 2485 | none | 5 | 23 | present (1) | absent (0) |
| CAS1-Delhi | 1343 | none | 1 | 23 | present (1) | absent (0) |
| CAS1-Delhi | 704 | none | 1 | 23 | present (1) | absent (0) |
| EAI2-Manila | 480 | none | 1 | 35 | present (1) | absent (0) |
| T4/T3 (WGS); none (HiSpoT) | none | 442 | 1 | 31 | present (1) | absent (0) |
| T1 (WGS); H3 (HiSpoT) | 121 | 122 | 1 | 31 | present (1) | absent (0) |

*Note:* In every discrepant isolate, the difference involved exactly one spacer, always detected as present by WGS and absent by HiSpoT. One CAS1-Kili isolate (SIT 21) and two EAI2-Manila isolates (SIT 19) gave identical spoligotypes by both methods and are not shown. Full isolate-level results are provided in Table S1.

25 out of 79 profiles (31%) showed discrepancies between the WGS and HiSpoT results, that is, about 70% agreement.

23 of 25 discrepancies were of 1 spacer difference, spacer present in WGS, absent in PCR-based HiSpoT targeted NGS.

2 of the 25 discrepancies (in samples 653-21 and 654-21) were due to extremely low counts for very weak detection in WGS, but strong good detection in HiSpoT. This unusual anomaly of spacers absent in WGS, but present in HiSpoT flagged the need to repeat the WGS NGS sequencing.

Our 79 isolates were taken from the years 2020 and 2021, years in which Israel had a significant large proportion of CAS family isolates (Figure 4). Most discrepancies were found within the CAS family, sub-group CAS1-Delhi SIT designations 25 and 2485, and one each of SIT designation 704 and 1343, all with the same one spacer difference, spacer 23 present in WGS, but absent in HiSpoT. All the WGS spoligotypes were assigned family “none” and SIT “none,” while all the HiSpoT spoligotypes were assigned the relevant family CAS and appropriate SIT.

One isolate, 590-20, showed a one spacer, spacer 35, difference (WGS present, HiSpoT absent) such that the WGS spoligotype was assigned “none” for family and SIT, while the HiSpoT spoligotype was assigned the relevant family EAI2-Manila and the appropriate SIT 480.

Two isolates, 264-21 and 656-21, showed a one spacer, spacer 31, difference (WGS present, HiSpoT absent) resulting in different family assignments for WGS versus HiSpoT derived spoligotypes.

### Confirmation and explanation of the one spacer discrepancies between WGS and HiSpoT derived spoligotypes

To resolve the genetic mechanism responsible for these single-spacer differences, sequence alignments were examined at the primer binding regions. As shown in Figure 9, and in Figure SI_5, all the CAS discrepancies concerning spacer 23 were explained by the same insertion element disruption of the primer site for mDRa, thus preventing PCR-based amplification (like in HiSpoT) of spacer 23 which was therefore absent on HiSpoT, but detected on WGS sequencing. The insertion element was labeled IS987, but due to an historical accident, the closely related IS6110 is often designated by ISfinder software as IS987, and indeed both belong to the same IS3 family of mobile element transposases. Figure SI_5 shows that other areas of insertions disrupted or even eliminated the primer sites for PCR amplification of spacer 31 and spacer 35.

**FIG 9.**
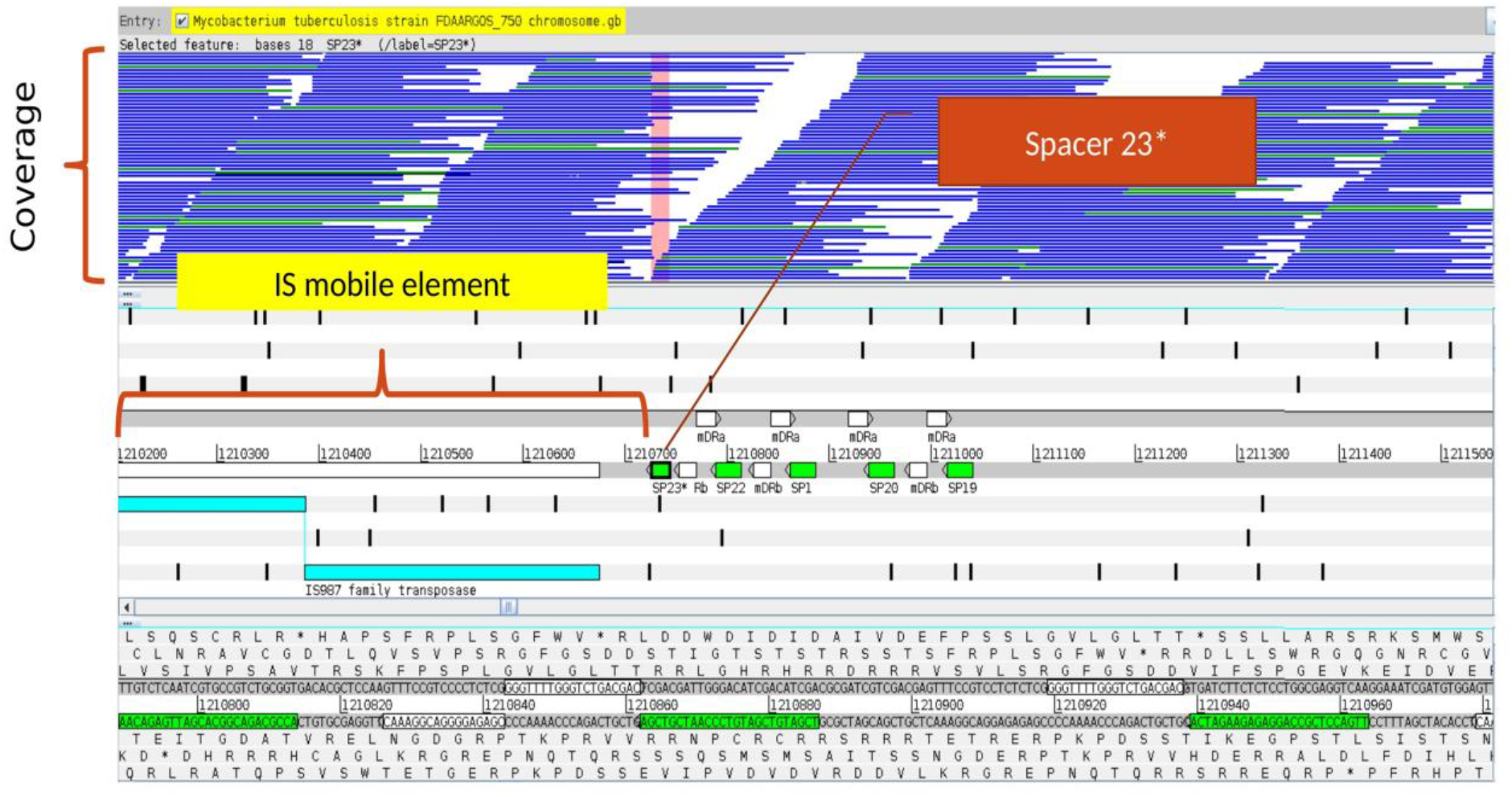
Molecular basis of the spacer 23 discrepancy between whole-genome sequencing (WGS) and HiSpoT in CAS1-Delhi isolates (example: SIT 2485). Read-coverage tracks (top) and the underlying reference sequence (bottom) show an IS*6110*/IS*987*-family insertion-sequence (IS3 family transposase) element inserted immediately adjacent to spacer 23 and its flanking mDRa primer-binding site. This insertion disrupts the PCR primer target, preventing amplification and detection of spacer 23 by HiSpoT, while WGS, which does not depend on this primer site, still detects the spacer. The figure was generated using the Artemis program (Sanger Institute). The same mechanism explains the spacer 23 discrepancies observed in SIT 25, 704, and 1343.

## DISCUSSION

### Specific features and benefits of the HiSpoT method

The HiSpoT novel one-tube dual barcode strategy to produce a ready-to-use targeted NGS amplicon library, gives highly accurate cost-effective (about 5-16% of the cost of WGS per sample, currently about 8 vs 50-150 US dollars), time-saving, labor-saving, high-throughput MTBC spoligotyping. Our method can be adapted easily to the spoligotyping of other microorganisms, for metagenomics applications, and in general to produce one-tube ready-to-use amplicon libraries for Illumina NGS use.

During PCR preparation the user can tell via real-time visualized melting curves whether the sample is good (MTBC), or needs repeating because it showed as negative or NTM (non-tuberculosis mycobacteria) (Fig. 5). Since the DR sequences are unique to the MTBC, a mixed culture of NTM and MTBC is expected to give a clear spoligotype in contrast to problematic WGS results. HiSpoT amplicon library preparation is a one tube dual-PCR affair - it does not require normalization, does not require a separate step for library preparation thus avoiding the costly and time consuming NGS library preparation kit, and requires the addition of only 1% phiX DNA versus the recommended 40% for standard Illumina NGS protocols (Recommendations for sequencing Illumina 16S libraries). Our novel method avoids the danger of cross contamination encountered in two step protocols which involve transfer of liquids containing millions of amplicons. Although in the studies reported in this paper, WGS quality DNA was used, HiSpoT could potentially use crude (heat extracted) DNA from MTBC cultures, and theoretically could be used successfully with crude DNA from sputum or other clinical submissions. HiSpoT sensitively detects nanograms of MTBC DNA, whereas WGS requires more MTBC DNA. In this regard, HiSpoT would be an excellent quality control (QC) procedure for WGS while providing first tier epidemiologic information: 1) negative WGS but positive HiSpoT results would flag the need to repeat WGS, 2) HiSpoT spoligotype MTBC family lineage inconsistent with WGS single nucleotide polymorphism (SNP) family lineage (14) would flag possible incorrect clinical sampling and the need to retest the clinical sample both by WGS and HiSpoT to confirm correct sampling by lineage congruence. In our work, there was perfect concordance between HiSpoT spoligotype family lineage and SNP family lineage (Table SI_6). Other groups have considered targeted NGS for spoligotyping MTBC, but the HiSpoT method is unique in eliminating the need for the time-consuming, labor-intensive, costly commercial library preparation kit (38, 39, 40, 41, 42).

### Sources of spoligotype polymorphism

Sources of spoligotype polymorphism include sequence polymorphism within a spacer, sequence polymorphism within DR sequences, and insertion element perturbation of primer targeted sequences. Experience to date suggests that insertion element derived WGS versus PCR spoligotyping based discrepancies usually affect only one spacer thus leaving a spoligotype that is consistent with LSP and SNP lineage identification. This is surprising given that insertion element presence in the genome has a rapid molecular clock in comparison to spoligotype polymorphism, for example observe the many IS6110 restriction fragment length polymorphisms (rflp) associated with a given CAS spoligotype (31). In particular, the DR region was early on identified as a hotspot for IS6110 insertion (43, 44).

IS6110 mobile element (aka transposase) insertion derived single spacer discrepancies between a spacer present by WGS, but absent by PCR based method (such as HiSpoT), have been extensively reported including for spacers 6, 9, 10, 11, 20, 24, 26, 31, 32, and 38, while spacer 18 discrepancy was attributed to spacer sequence variations and spacer 42 discrepancy was attributed to a mutated DR sequence (45, 46, 47, 48). It is clear that in order to maintain connection with the massive PCR based spoligotype databases that provide biogeographic information and other attributes, HiSpoT PCR based targeted NGS is a low cost alternative or simultaneous option to WGS for quick high throughput generation of informative (PCR based) spoligotypes. There are also quality control aspects to performing HiSpoT NGS along with WGS NGS.

### Software for analysis of HiSpoT or WGS NGS Illumina fastq output

A number of software solutions have been offered for analysis of PCR based and WGS reads and genome sequences for MTBC spoligotypes (35, 49, 50, 51, 52, 53, 54, 55, 56, 57, 58, 59, 60). Our software was independently developed and validated, and to the best of our knowledge utilizes efficient unique algorithms that are especially fast in analyzing HiSpoT Illumina paired-end fastq reads output. The Spoligotyping_MTBC_gcl.ipynb software, or its VIBE version HiSpoT_Spoligotyping_v8_ready.ipynb, takes about 30 seconds to analyze HiSpoT output for 96 samples.

In summary, HiSpoT is a one-step dual-barcode PCR-based targeted NGS, and is a simpler, faster, cheaper, more high-throughput spoligotyping method than extant PCR-based or WGS methods. An especially attractive benefit is that HiSpoT eliminates the need for the labor-intensive, time-consuming, costly commercial library preparation kit. HiSpoT is easily performed on NGS platforms that are widely available even in resource challenged countries, and de-multiplexing is performed automatically by the sequencer (like MiSeq). The HiSpoT targeted NGS results are bioinformatically analyzed in seconds to a few minutes by software which reports spoligotypes, SITs, and families. The spoligotyping softwares are available as interactive python notebook (ipynb) files with accompanying modules and validation data from GitHub, and easily deployed as Google Colab Jupyter Notebooks with the accompanying files in Google Drive.

## ACKNOWLEDGMENTS

We thank the staff of the National Public Health Laboratory, Ministry of Health, Tel Aviv, Israel, for technical assistance with sample processing and sequencing.

## Funding

[Author to insert: state all funding sources and grant numbers, or include the statement: “This research received no specific grant from any funding agency in the public, commercial, or not-for-profit sectors.”]

## Author contributions

Paul J. Freidlin: software concept, development, and validation; molecular biology of MTBC; data analysis (Table 1 and Table S1); writing - original draft and review of subsequent drafts. Israel Nissan: HiSpoT design and development; project coordination; preparation of the Google Colab VIBE version; bioinformatic analysis; writing - review and editing. Ephraim Fass, Mor Rubinstein, Inna Kutikov, Gal Zizelski Valenci, Yelena Losev, Zeev Dveyrin, and Efrat Rorman contributed to laboratory work, data generation, and/or manuscript review.

## Conflicts of interest

The HiSpoT method is described in an open-access, royalty-free patent (https://patents.google.com/patent/US20210355526A1/en). The authors declare no other competing interests.

## DATA AVAILABILITY

The Spoligotyping_MTBC_gcl and HiSpoT_Spoligotyping_v8_ready software packages (Google Colaboratory/Jupyter notebooks, supporting Python modules, and validation fastq dataset and results) are freely available for download from GitHub (https://github.com/isrulik67/HiSpoT-dual-barcode-NGS-spoligotyping-of-M.-tuberculosis/releases/tag/v8.1) and from zenodo. Org (DOI: 10.5281/zenodo.23948119). The HiSpoT one-tube dual-barcode PCR method is described in open-access patent US20210355526A1 (https://patents.google.com/patent/US20210355526A1/en). Detailed primer sequences, barcodes, and assay protocols for up to 96 samples are provided in Supplementary Information (Tables S1-S3, Table S4, and Fig. S5); underlying isolate-level spoligotype comparison data are provided in Table S1. The whole-genome sequencing read set for isolate T3_0525-11 (spoligotyping result provided) was previously deposited in GenBank for public access (Freidlin et al., reference 22 in this manuscript).

## Notes

https://doi.org/10.5281/zenodo.23948119

https://github.com/isrulik67/HiSpoT-dual-barcode-NGS-spoligotyping-of-M.-tuberculosis/tree/v8.1

### Competing Interest Statement

The authors have declared no competing interest.

## REFERENCES

1. World Health Organization. Global tuberculosis report 2025. Geneva: World Health Organization; 2025. https://www.who.int/publications/i/item/9789240116924

2. Filliol I, Driscoll JR, Van Soolingen D, Kreiswirth BN, Kremer K, Valetudie G, Anh DD, Barlow R, Banerjee D, Bifani PJ, et al. Global distribution of Mycobacterium tuberculosis spoligotypes. Emerg Infect Dis. 2002;8(11):1347–1349. 10.3201/eid0811.020125

3. Cohen KA, Abeel T, Manson McGuire A, Desjardins CA, Munsamy V, Shea TP, et al. Evolution of extensively drug-resistant tuberculosis over four decades. PLoS Med. 2015;12:e1001880. 10.1371/journal.pmed.1001880

4. Filliol I, Driscoll JR, van Soolingen D, Kreiswirth BN, Kremer K, Valétudie G, et al. Snapshot of moving and expanding clones of Mycobacterium tuberculosis assessed by spoligotyping. J Clin Microbiol. 2003;41:1963–1970. 10.1128/JCM.41.5.1963-1970.2003

5. Goldblatt D, Rorman E, Chemtob D, Freidlin PJ, Cedar N, Kaidar-Shwartz H, Dveyrin Z, Mor Z. Molecular epidemiology and mapping of tuberculosis in Israel: do migrants transmit the disease to locals? Int J Tuberc Lung Dis. 2014 Sep;18(9):1085–91. 10.5588/ijtld.14.0186

6. Brudey K, Driscoll JR, Rigouts L, Prodinger WM, Gori A, Al-Hajoj SA, et al. Mycobacterium tuberculosis complex genetic diversity: mining the fourth international spoligotyping database (SpolDB4) for classification, population genetics and epidemiology. BMC Microbiol. 2006;6:23. 10.1186/1471-2180-6-23

7. Couvin D, Allaguy AS, Ez-Zari A, Jagielski T, Rastogi N. Molecular typing of Mycobacterium tuberculosis: a review of current methods, databases, softwares, and analytical tools. FEMS Microbiol Rev. 2025;49:fuaf017. 10.1093/femsre/fuaf017

8. Demay C, Liens B, Burguière T, Hill V, Couvin D, Millet J, et al. SITVITWEB: a publicly available international multimarker database for studying Mycobacterium tuberculosis genetic diversity. Infect Genet Evol. 2012;12:755–766. 10.1016/j.meegid.2012.02.004

9. Ishino Y, Shinagawa H, Makino K, Amemura M, Nakata A. Nucleotide sequence of the iap gene, responsible for alkaline phosphatase isozyme conversion in Escherichia coli, and identification of the gene product. J Bacteriol. 1987;169(12):5429–5433. 10.1128/jb.169.12.5429-5433.1987

10. Groenen PM, Bunschoten AE, van Soolingen D, van Embden JD. Nature of DNA polymorphism in the direct repeat cluster of Mycobacterium tuberculosis. Mol Microbiol. 1993;10:1057–1065. 10.1111/j.1365-2958.1993.tb00976.x

11. Kamerbeek J, Schouls L, Kolk A, van Agterveld M, van Soolingen D, Kuijper S, et al. Simultaneous detection and strain differentiation of Mycobacterium tuberculosis. J Clin Microbiol. 1997;35:907–914. 10.1128/jcm.35.4.907-914.1997

12. van Embden JD, van Gorkom T, Kremer K, Jansen R, van Der Zeijst BA, Schouls LM. Genetic variation and evolutionary origin of the direct repeat locus of Mycobacterium tuberculosis complex bacteria. J Bacteriol. 2000 May;182(9):2393–401. https://doi.org/10.1128/JB,182.9.2393-2401.2000

13. van Soolingen D, Qian L, de Haas PE, Douglas JT, Traore H, Portaels F, et al. Predominance of a single genotype in East Asia. J Clin Microbiol. 1995;33:3234– 3238. 10.1128/jcm.33.12.3234-3238.1995

14. Coll F, McNerney R, Guerra-Assunção JA, Glynn JR, Perdigão J, Viveiros M, et al. A robust SNP barcode for typing Mycobacterium tuberculosis complex strains. Nat Commun. 2014;5:4812. 10.1038/ncomms5812

15. Freidlin P, Goldblatt D, Kaidar-Shwartz H, Dveyrin Z, Rorman E. Quality assurance for molecular epidemiology of tuberculosis methods in the mycobacterium reference laboratory. Accred Qual Assur. 2011;16(12):623–635. 10.1007/s00769-011-0819-5

16. Popovici O, Monk P, Chemtob D, Chiotan D, Freidlin PJ, Groenheit R, Haanperä M, Homorodean D, Mansjö M, Robinson E, Rorman E, Smith G, Soini H, Van Der Werf MJ. Cross-border outbreak of extensively drug-resistant tuberculosis linked to a university in Romania. Epidemiol Infect. 2018 May;146(7):824–831. 10.1017/S095026881800047X

17. Abadia E, Zhang J, Ritacco V, Kremer K, Ruimy R, Rigouts L, et al. The use of microbead-based spoligotyping for Mycobacterium tuberculosis complex to evaluate the quality of the conventional method: providing guidelines for quality assurance when working on membranes. BMC Infect Dis. 2011;11:110. 10.1186/1471-2334-11-110

18. Bespyatykh JA, Zimenkov DV, Shitikov EA, Kulagina EV, Lapa SA, Gryadunov DA, et al. Spoligotyping of Mycobacterium tuberculosis complex isolates using hydrogel oligonucleotide microarrays. Infect Genet Evol. 2014;26:41–46. 10.1016/j.meegid.2014.04.024

19. Honisch C, Mosko M, Arnold C, Gharbia SE, Diel R, Niemann S. Replacing reverse line blot hybridization spoligotyping. J Clin Microbiol. 2010;48:1520–1526. 10.1128/JCM.02299-09

20. Sola C, Abadia E, Le Hello S, Weill FX. High-throughput CRISPR typing of Mycobacterium tuberculosis complex. Methods Mol Biol. 2015;1311:39–53. 10.1007/978-1-4939-2687-9_6

21. Xia Z, Su B, Tu C, Sun S, Tan Y, Xu Y, Li Q. Single-tube culture-independent spoligotyping protocol. J Clin Microbiol. 2024;62:e01183–23. 10.1128/jcm.01183-23

22. Freidlin PJ, Nissan I, Luria A, Goldblatt D, Schaffer L, Kaidar-Shwartz H, et al. Structure and variation of CRISPR and CRISPR-flanking regions in Mycobacterium tuberculosis complex strains. BMC Genomics. 2017;18:168. 10.1186/s12864-017-3560-6

23. Gagneux S, DeRiemer K, Van T, Kato-Maeda M, de Jong BC, Narayanan S, et al. Variable host–pathogen compatibility in Mycobacterium tuberculosis. Proc Natl Acad Sci USA. 2006;103:2869–2873. 10.1073/pnas.0511240103

24. Kato-Maeda M, Gagneux S, Flores LL, Kim EY, Small PM, Desmond EP, Hopewell PC. Strain classification of Mycobacterium tuberculosis. Int J Tuberc Lung Dis. 2011;15:131–133. No doi available

25. Tsolaki AG, Hirsh AE, DeRiemer K, Enciso JA, Wong MZ, Hannan M, Goguet de la Salmoniere YO, Aman K, Kato-Maeda M, Small PM. Functional and evolutionary genomics of Mycobacterium tuberculosis: insights from genomic deletions in 100 strains. Proc Natl Acad Sci U S A. 2004;101(14):4865–4870. 10.1073/pnas.0305634101

26. Vasconcellos SE, Acosta CC, Gomes LL, Conceição EC, Lima KV, de Araujo MI, et al. Strain classification of Mycobacterium tuberculosis in Brazil. PLoS One. 2014;9:e107747. 10.1371/journal.pone.0107747

27. Garneau JE, Dupuis ME, Villion M, Romero DA, Barrangou R, Boyaval P, Fremaux C, Horvath P, Magadan AH, Moineau S. The CRISPR/Cas bacterial immune system cleaves bacteriophage and plasmid DNA. Nature. 2010;468(7320):67–71. 10.1038/nature09523

28. Makarova KS, Shmakov SA, Wolf YI, Mutz P, Altae-Tran H, Beisel CL, et al. Evolutionary classification of CRISPR-Cas systems. Nat Microbiol. 2025;10:3346– 3361. 10.1038/s41564-025-02180-8

29. Marraffini LA, Sontheimer EJ. CRISPR interference limits horizontal gene transfer in staphylococci by targeting DNA. Science. 2008;322(5909):1843–1845. 10.1126/science.1165771

30. Mojica FJ, Rodriguez-Valera F. The discovery of CRISPR in archaea and bacteria. FEBS J. 2016;283:3162–3169. 10.1111/febs.13766

31. Freidlin PJ, Goldblatt D, Kaidar-Shwartz H, Rorman E. Polymorphic exact tandem repeat A (PETRA): a newly defined lineage of mycobacterium tuberculosis in israel originating predominantly in Sub-Saharan Africa. J Clin Microbiol. 2009 Dec;47(12):4006–20. 10.1128/JCM.01270-09

32. Dale JW, Brittain D, Cataldi AA, Cousins D, Crawford JT, Driscoll J, Heersma H, Lillebaek T, Quitugua T, Rastogi N, et al. Spacer oligonucleotide typing of bacteria of the Mycobacterium tuberculosis complex: recommendations for standardised nomenclature. Int J Tuberc Lung Dis. 2001;5(3):216–219

33. Gessese AT, Kinde MZ, Eshetu T, Kerisew B. Whole-genome sequencing analysis of Mycobacterium tuberculosis isolates from Ethiopia. Sci Rep. 2025;15:16076. 10.1038/s41598-025-01241-6

34. Horan, K. A., Viberg, L., Ballard, S. A., Globan, M., Wirth, W., Bond, K., Webb, J. R., Dorji, T., Williamson, D. A., Sait, M. L., Tay, E. L., Denholm, J. T., Howden, B. P., Seemann, T., & Sherry, N. L. (2025). Bringing tuberculosis genomics to the clinic: development and validation of a comprehensive pipeline to predict antimicrobial susceptibility from genomic data, accredited to ISO standards. The Lancet Digital Health, 7(12), 100939. 10.1016/j.landig.2025.100939

35. Gomgnimbou MK, Hernández-Neuta I, Panaiotov S, Bachiyska E, Palomino JC, Martin A, del Portillo P, Refregier G, Sola C. Tuberculosis-spoligo-rifampin-isoniazid typing: an all-in-one assay technique for surveillance and control of multidrug-resistant tuberculosis on Luminex devices. J Clin Microbiol. 2013 Nov;51(11):3527–34. doi: 10.1128/JCM.01523-13.

36. Napier G, Couvin D, Refrégier G, Guyeux C, Meehan CJ, Sola C, et al. Comparison of in silico predicted spoligotypes from WGS data. Sci Rep. 2023;13:11368. 10.1038/s41598-023-38384-3

37. Xia E, Teo YY, Ong RT. SpoTyping: fast and accurate in silico spoligotyping. Genome Med. 2016;8:19. 10.1186/s13073-016-0270-7

38. Genestet C, Baffert Y, Vallée M, Bernard A, Benito Y, Lina G, et al. Development, evaluation, and implementation of targeted next-generation sequencing spoligotyping. Int J Mol Sci. 2022;23:11302. 10.3390/ijms231911302

39. Gou B, Zhang C, Song Z, Huang Z, Lu S. Targeted next-generation sequencing for rapid tuberculosis detection: a systematic review and meta-analysis. Clin Chim Acta. 2025 Sep 1;577:120469. 10.1016/j.cca.2025.120469

40. Hess JF, Kohl TA, Kotrová M, Rönsch K, Paprotka T, Mohr V, Hutzenlaub T, Brüggemann M, Zengerle R, Niemann S, Paust N. Library preparation for next generation sequencing: A review of automation strategies. Biotechnol Adv. 2020 Jul-Aug;41:107537. 10.1016/j.biotechadv.2020.107537

41. Mokrousov I, Chernyaeva E, Vyazovaya A, Sinkov V, Zhuravlev V, Narvskaya O. Next-generation sequencing of Mycobacterium tuberculosis. Emerg Infect Dis. 2016;22:1127–1129. 10.3201/eid2206.152051

42. Slatko BE, Gardner AF, Ausubel FM. Overview of Next-Generation Sequencing Technologies. Curr Protoc Mol Biol. 2018 Apr;122(1):e59. 10.1002/cpmb.59

43. Fang Z, Morrison N, Watt B, Doig C, Forbes KJ 1998. IS6110 Transposition and Evolutionary Scenario of the Direct Repeat Locus in a Group of Closely Related Mycobacterium tuberculosis Strains. J Bacteriol 1998; 180:

44. Sampson SL, Warren RM, Richardson M, Victor TC, Jordaan AM, van der Spuy GD, van Helden PD. IS6110-mediated deletion polymorphism in the direct repeat region of clinical isolates of Mycobacterium tuberculosis. J Bacteriol. 2003;185(9):2856–2866. 10.1128/JB.185.9.2856-2866.2003

45. Bakuła Z, Dziurzyński M, Decewicz P, Bakonytė D, Vasiliauskaitė L, Nakčerienė B, et al. Spoligotyping of Mycobacterium tuberculosis – comparing in vitro and in silico approaches. Infect Genet Evol. 2023;115:105508. 10.1016/j.meegid.2023.105508

46. Benjamin WH Jr, Lok KH, Harris R, Brook N, Bond L, Mulcahy D, Robinson N, Pruitt V, Kirkpatrick DP, Kimerling ME, Dunlap NE. Identification of a contaminating Mycobacterium tuberculosis strain with a transposition of an IS6110 insertion element resulting in an altered spoligotype. J Clin Microbiol. 2001 Mar;39(3):1092–6. doi: 10.1128/JCM.37.3.1092-1096.2001

47. Genestet C, Hodille E, Bernard A, Vallée M, Lina G, Le Meur A, et al. Consistency of Mycobacterium tuberculosis complex spoligotyping between membrane-based and in silico approaches. Microbiol Spectr. 2022;10:e0022322. 10.1128/spectrum.00223-22

48. Lok, K. H., Benjamin, W. H., Kimerling, M. E., Pruitt, V., Mulcahy, D., Robinson, N.…Dunlap, N. E. (2002). Molecular Typing of Mycobacterium tuberculosis Strains with a Common Two-Band IS6110 Pattern. Emerging Infectious Diseases, 8(11), 1303–1305. 10.3201/eid0811.020292

49. Coll F, Mallard K, Preston MD, Bentley S, Parkhill J, McNerney R, Martin N, Clark TG. SpolPred: rapid and accurate prediction of Mycobacterium tuberculosis spoligotypes from short genomic sequences. Bioinformatics. 2012;28:2991–2993. 10.1093/bioinformatics/bts544

50. Couvin D, Segretier W, Stattner E, Rastogi N. Novel methods included in SpolLineages tool for fast and precise prediction of Mycobacterium tuberculosis complex spoligotype families. Database (Oxford). 2020;2020:baaa108. 10.1093/database/baaa108

51. Couvin D, Stattner E, Segretier W, Cazenave D, Rastogi N. simpiTB: a pipeline designed to extract meaningful information from whole-genome sequencing data of Mycobacterium tuberculosis complex. Infect Genet Evol. 2023;113:105466. 10.1016/j.meegid.2023.105466

52. Crawford JT, Braden CR, Schable BA, Onorato IM. National Tuberculosis Genotyping and Surveillance Network: design and methods. Emerg Infect Dis. 2002;8:1192–1196. 10.3201/eid0811.020296

53. Hijikata M, Keicho N, Duc LV, Maeda S, Hang NTL, Matsushita I, Kato S. Spoligotyping and whole-genome sequencing of lineage 1 strains in Vietnam. PLoS One. 2017;12:e0186800. 10.1371/journal.pone.0186800

54. Kohl TA, Diel R, Harmsen D, Rothgänger J, Walter KM, Merker M, et al. Whole-genome-based Mycobacterium tuberculosis surveillance. J Clin Microbiol. 2014;52:2479–2486. 10.1128/JCM.00567-14

55. Muzyukina P, Soutourina O. CRISPR genotyping methods: from spoligotyping to machine learning. Biochimie. 2024;217:66–73. 10.1016/j.biochi.2023.07.017

56. Rasoahanitralisoa R, Rakotosamimanana N, Stucki D, Sola C, Gagneux S, Rasolofo Razanamparany V. Evaluation of spoligotyping and SNPs in Madagascar. PLoS One. 2017;12:e0186088. 10.1371/journal.pone.0186088

57. Sekizuka T, Yamashita A, Murase Y, Iwamoto T, Mitarai S, Kato S, Kuroda M. TGS-TB: total genotyping solution. PLoS One. 2015;10:e0142951. 10.1371/journal.pone.0142951

58. Shabbeer A, Cowan LS, Ozcaglar C, Rastogi N, Vandenberg SL, Yener B, Bennett KP. TB-Lineage: an online classification tool. Infect Genet Evol. 2012;12:789–797. 10.1016/j.meegid.2012.02.010

59. Tang C, Reyes JF, Luciani F, Francis AR, Tanaka MM. spolTools: online utilities for analyzing spoligotypes. Bioinformatics. 2008;24:2414–2415. 10.1093/bioinformatics/btn434

60. Weniger T, Krawczyk J, Supply P, Harmsen D, Niemann S. Online tools for polyphasic analysis of Mycobacterium tuberculosis. Infect Genet Evol. 2012;12:748–754. 10.1016/j.meegid.2012.01.021

