## Supplemental Data 1 for "HiSpoT: a novel one-tube dual-barcode PCR method for NGS spoligotyping amplicon library preparation in *Mycobacterium tuberculosis* complex"

In isolate 609-20, CAS1-Delhi SIT 2485, the mDRa primer for Spacer 23 is rendered inoperable by insertion of a mobile element that prevents the PCR amplification of Spacer 23.

```
1210441  CCCGCCGCAA GCGCTTCAGC TCAGCGGATT CTTCGGTCGT GGTCCCGGGC CGTGCGCCGG CATCGACCTG CGCCTGGCGC ACCCACTTAC GCACCGTCTC CGCGCAGCCA ACACCAAGTA GACGGGCGAC CTCACTGATC
         GGGCGGCGTT CGCGAAGTCG AGTCGCCTAA GAAGCCAGCA CCAGGGCCCG GCACGCGGCC GTAGCTGGAC GCGGACCGCG TGGGTGAATG CGTGGCAGAG GCGCGTCGGT TGTGGTTCAT CTGCCCGCTG GAGTGACTAG

1210581  GCTGCCCACT CCGAATCGTG CTGACCGCGG ATCTCTGCGA CCATCCGCAC CGCCCGCTCA CGCAGCTCCG GCGGGTACCT CCTCGATGAA CCACCTGACA TGACCCCATC CTTTCCAAGA ACTGGAGTCT CCGGACATGC
         CGACGGGTGA GGCTTAGCAC GACTGGCGCC TAGAGACGCT GGTAGGCGTG GCGGGCGAGT GCGTCGAGGC CGCCCATGGA GGAGCTACTT GGTGGACTGT ACTGGGGTAG GAAAGGTTCT TGACCTCAGA GGCCTGTACG

                                                                                                                                        ◄ DRb
                                                                                                                                  CAAAGGCA GGGGAGAGCC
                                                   ◄ DRb
                                              C AAAGGCAGGG GAGAGCC
1210721  CGGGGCGGTT CAGACCGCAT CAGCGATGCT GAGCTGAGGG TTTCCGTCCC CTCTCGGGGT TTTGGGTCTG ACGACTTGTC TCAATCGTGC CGTCTGCGGT GACACGCTCC AAGTTTCCGT CCCCTCTCGG GGTTTTGGGT
         GCCCCGCCAA GTCTGGCGTA GTCGCTACGA CTCGACTCCC AAAGGCAGGG GAGAGCCCCA AAACCCAGAC TGCTGAACAG AGTTAGCACG GCAGACGCCA CTGTGCGAGG TTCAAAGGCA GGGGAGAGCC CCAAAACCCA
                    >>.....SP23mut.....>>
                                                             >>......DRa.......>>
                                                                               >>.........SP22.........>>
                                                                                                                                                   >>.DRa...>

                                                                                                                                                    ◄ DRb
                                                                                                                                              CAAAG GCAGGGGAGA
1210861  CTGACGACTC GACGATTGGG ACATCGACAT CGACGCGATC GTCGACGAGT TTCCGTCCTC TCTCGGGGTT TTGGGTCTGA CGACGTGATC TTCTCTCCTG GCGAGGTCAA GGAAATCGAT GTGGAGTTTC CGTCCCCTCT
         GACTGCTGAG CTGCTAACCC TGTAGCTGTA GCTGCGCTAG CAGCTGCTCA AAGGCAGGAG AGAGCCCCAA AACCCAGACT GCTGCACTAG AAGAGAGGAC CGCTCCAGTT CCTTTAGCTA CACCTCAAAG GCAGGGGAGA
         >.DRa.>>
                 >>.........SP21.........>>
                                                                     >>......DRa.......>>
                                                                                          >>.........SP20.........>>

         DRb
         GCC
1211001  CGGGGTTTTG GGTCTGACGA CCACCAGGAT CAGCGCCAAG CCAGTTAGCG CAATCCAGTT TCCGTCCCCT CTCGGGGTTT TGGGTCTGAC GACCTCCCGG ACCATCTGCA GCTCGCCCGG GTCCATGCGG TTTCCGTCCC
         GCCCCAAAAC CCAGACTGCT GGTGGTCCTA GTCGCGGTTC GGTCAATCGC GTTAGGTCAA AGGCAGGGGA GAGCCCCAAA ACCCAGACTG CTGGAGGGCC TGGTAGACGT CGAGCGGGCC CAGGTACGCC AAAGGCAGGG
            >>......DRa.......>>
                              >>.........SP19.........>>
                                                                                                                >>.........SP18.........>>
```

### *M. tuberculosis* 264-21map with H37Rv

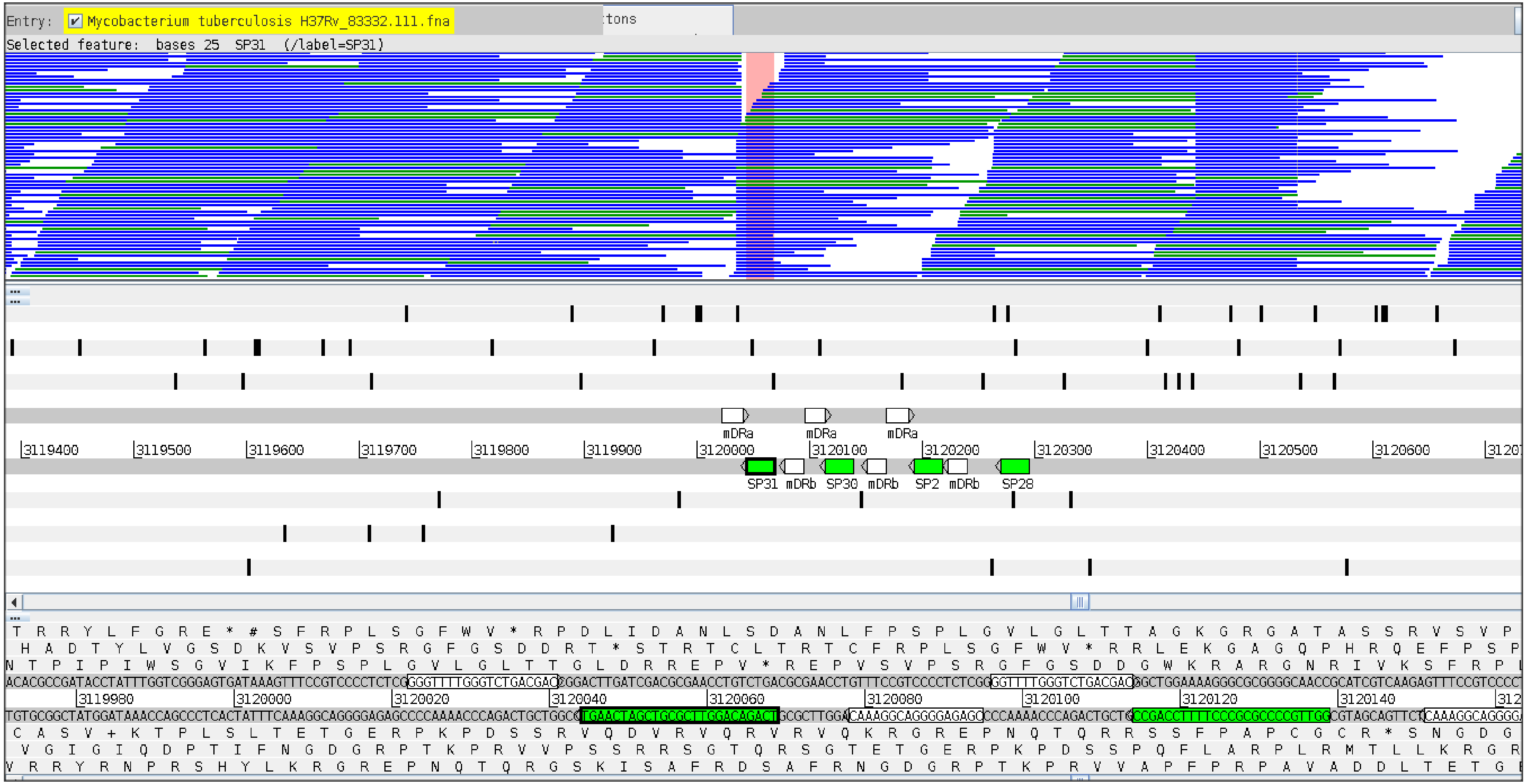


The reads are disrupted within the mDRa preventing amplification of spacer 31.

### *M. tuberculosis*_264-21 *de-novo* assembly

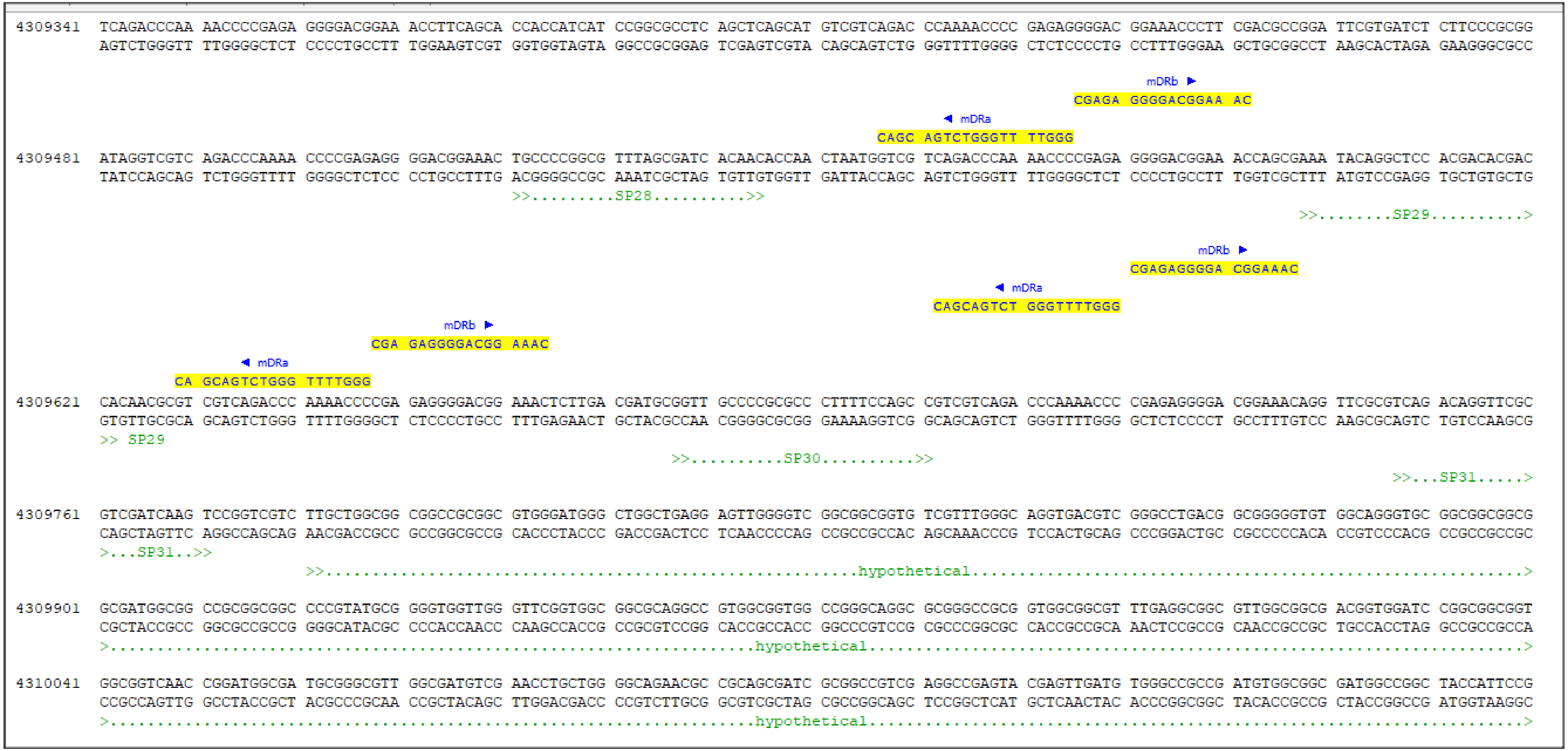


In the middle of the mDRa sequence there is a big insert of a mobile element that disrupts PCR amplification of spacer 31.

#### *Mycobacterium tuberculosis* 590-20, EAI2-manila, SIT 480

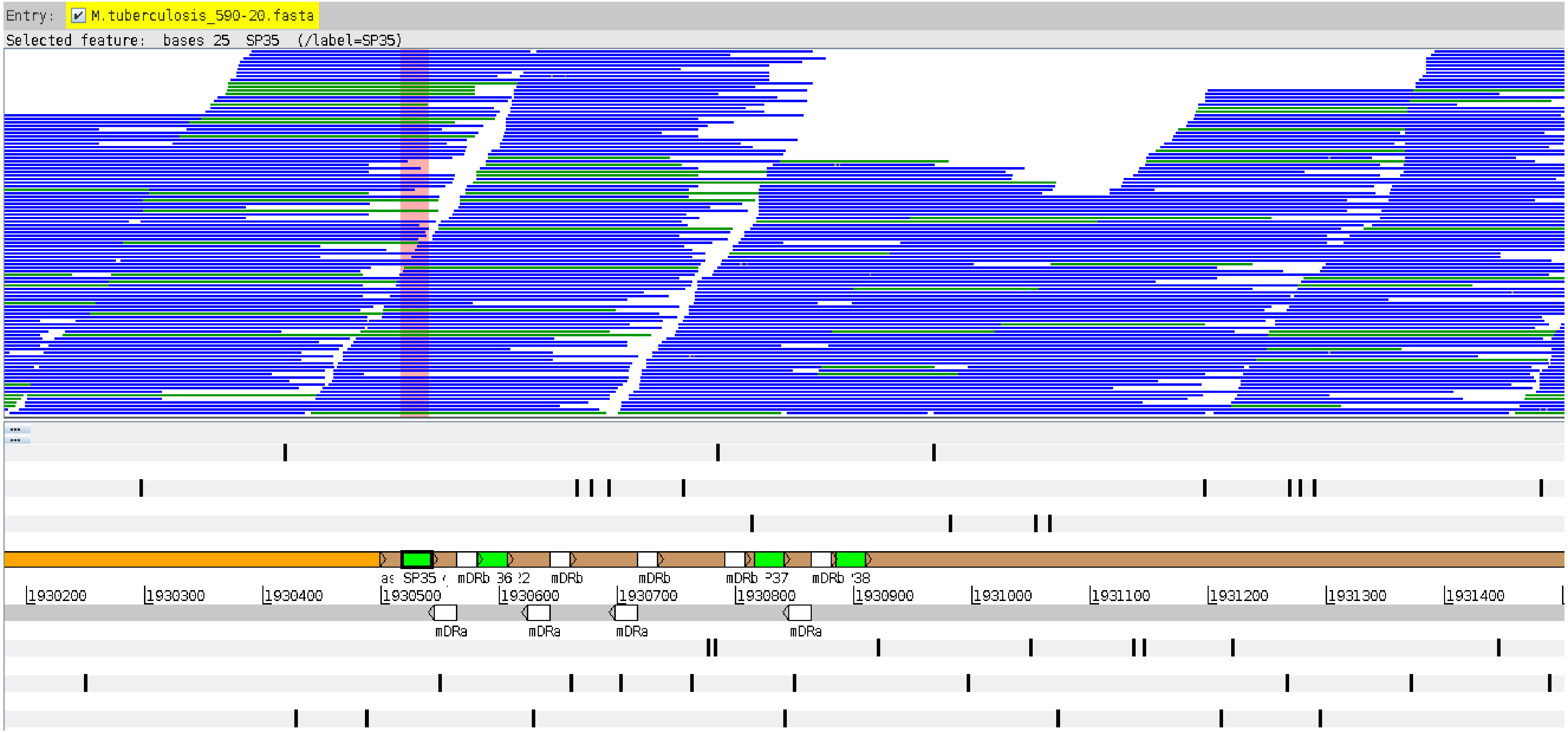


Several genes, perhaps transferred by a mobile element, have seemingly replaced mDRb thus preventing amplification of spacer 35.

### *Mycobacterium tuberculosis* 590-20, EAI2-manila, SIT 480

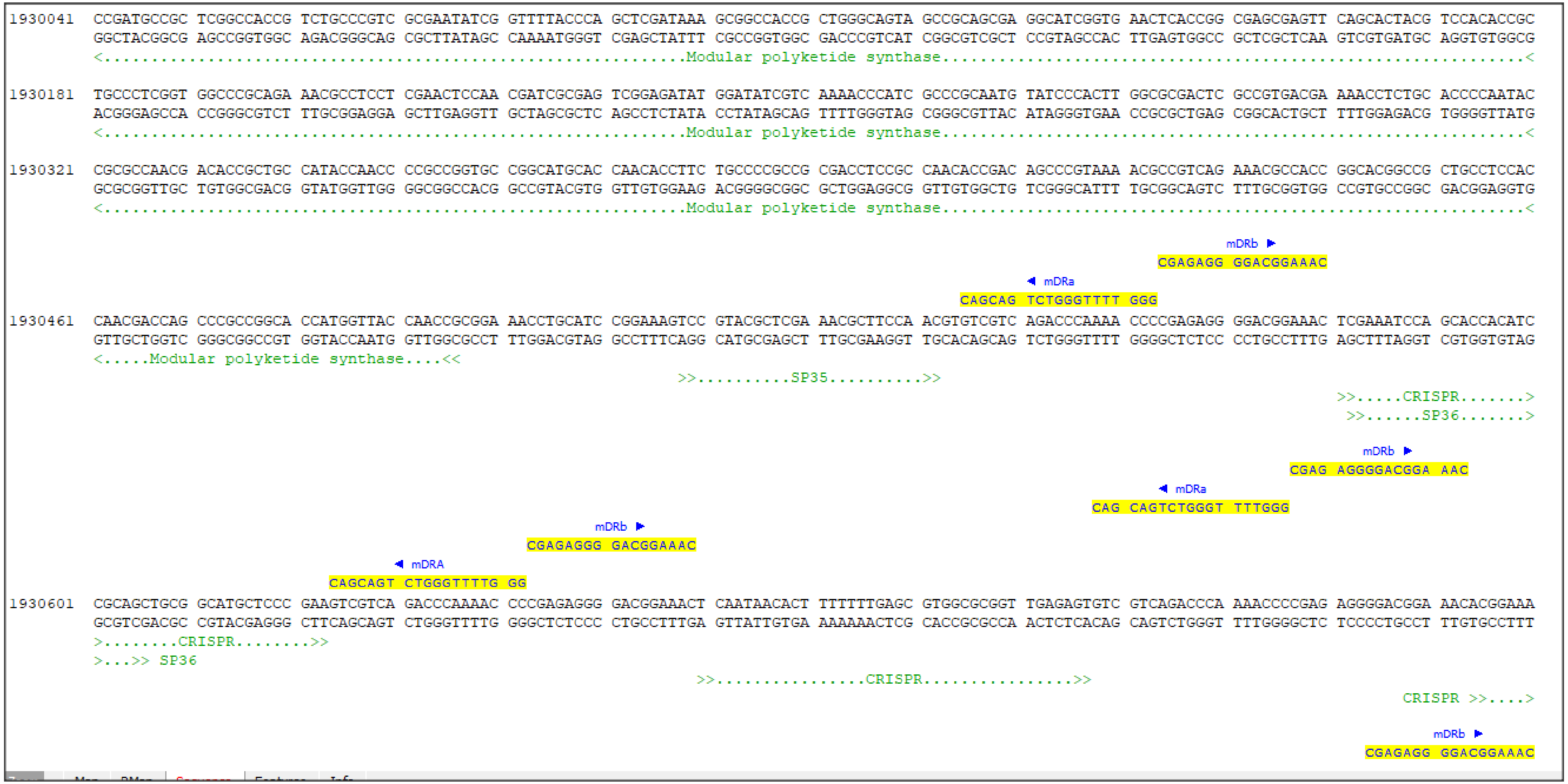


Several genes, perhaps transferred by a mobile element, have seemingly replaced mDRb thus preventing amplification of spacer 35. The genes belong to the PKS family.
