## supplementary information SI_2 for "HiSpoT: a novel one-tube dual-barcode PCR method for NGS spoligotyping amplicon library preparation in *Mycobacterium tuberculosis* complex"

**Appendix SOP: HiSpoT wet-lab protocol for MiSeq-compatible MTBC spoligotyping**

Version: draft 1.2

### 1. Purpose

To describe the wet-lab HiSpoT procedure for high-throughput sequencing-based spoligotyping of MTBC DNA, including primer preparation, 96-well reaction setup, PCR, pooling, library cleanup, library dilution, MiSeq sample sheet preparation, sequencing setup, and run-level QC records.

### 2. Scope and biosafety

The procedure applies to purified DNA from MTBC isolates after validated inactivation/extraction. All handling of live MTBC must follow institutional BSL-3 procedures. This SOP begins at the purified DNA stage. Negative and positive controls must be included in every plate.

### 3. Principle of the method

HiSpoT targets the classical spoligotyping DR/spacer locus. Universal DR primers and indexed primers generate amplicons in a closed tube, using std-PCR chemistry, that then are sequenced on a MiSeq. The software determines the presence or absence of the 43 classical spacers and reports the corresponding binary profile, octal spoligotype, SIT and family assignment.

### 4. Required equipment and materials

- Purified/inactivated MTBC DNA samples and nuclease-free water negative controls.
- KOD Hot Start DNA Polymerase and supplied 10× buffer.
- 2 mM dNTP mix; 25 mM MgCl₂; nuclease-free water.
- LightCycler 480 ResoLight dye, 20×, or equivalent dye used in the validated assay.
- HiSpoT primer stocks as listed in Section 6.
- 96-well PCR plate, optical or high-quality sealing film, strip tubes or primer reservoir plate.
- Calibrated single- and multichannel pipettes; preferably electronic repeat pipette for master mix dispensing.
- PCR instrument capable of the cycling program below; the real run used CFX96/CFX Manager.
- Promega ProNex cleanup kit or validated equivalent; magnetic rack.
- DeNovix/Qubit or equivalent dsDNA quantification; fragment analyzer/Bioanalyzer/TapeStation or equivalent smear analysis.
- Illumina MiSeq and compatible MiSeq reagent kit supporting 2×151 reads.

### 5. Workflow overview

| **Step** | **Action** |
| --- | --- |
| 1 | Prepare primer stocks and DR primer mix. |
| 2 | Enter sample IDs in the 96-well Excel reaction-planning template; export the MiSeq sample sheet. |
| 3 | Dispense indexed reverse primers by row and indexed forward primers by column. |
| 4 | Prepare common master mix excluding the indexed primers and DNA; dispense 14 µL per well. |
| 5 | Add 2 µL DNA template per sample; seal and run PCR. |
| 6 | Pool 10 µL from each well, clean the pooled library, quantify and check size distribution. |
| 7 | Normalize/denature/dilute the pooled library, prepare MiSeq sample sheet, and sequence. |
| 8 | Transfer FASTQ files to the analysis workflow and review QC output. |

### 6. Primer order list

Order primers desalted or HPLC-purified according to laboratory preference.

6A: DR specific primers.

| **Name** | **Direction** | **Sequence** | **Length** | **Notes** |
| --- | --- | --- | --- | --- |
| UT-mDRaSP15 | forward | TCGTCGGCAGCGTCAGATGTGTATAAGAGACAGGGTTTTGGGTCTGATCCG | 51 | In order to improve spacer 15 amplification |
| UT-mDRa1 | forward | TCGTCGGCAGCGTCAGATGTGTATAAGAGACAGGGGTTTTGGGTCTGACGAC | 53 | Designed in-house for this study |
| UT-mDRa2 | forward | TCGTCGGCAGCGTCAGATGTGTATAAGAGACAGNGGGTTTTGGGTCTGACGAC | 54 | Designed in-house for this study |
| UT-mDRa3 | forward | TCGTCGGCAGCGTCAGATGTGTATAAGAGACAGNNGGGTTTTGGGTCTGACGAC | 55 | Designed in-house for this study |
| UT-mDRa4 | forward | TCGTCGGCAGCGTCAGATGTGTATAAGAGACAGNNNGGGTTTTGGGTCTGACGAC | 56 | Designed in-house for this study |
| UT-mDRb1 | Reverse | GTCTCGTGGGCTCGGAGATGTGTATAAGAGACAGCGAGAGGGGACGGAAAC | 51 | Designed in-house for this study |
| UT-mDRb2 | Reverse | GTCTCGTGGGCTCGGAGATGTGTATAAGAGACAGNCGAGAGGGGACGGAAAC | 52 | Designed in-house for this study |
| UT-mDRb3 | Reverse | GTCTCGTGGGCTCGGAGATGTGTATAAGAGACAGNNCGAGAGGGGACGGAAAC | 53 | Designed in-house for this study |
| UT-mDRb4 | Reverse | GTCTCGTGGGCTCGGAGATGTGTATAAGAGACAGNNNCGAGAGGGGACGGAAAC | 54 | Designed in-house for this study |

6B: Flow cell adaptors and dual barcode primers.

| **Name** | **Direction** | **Sequence** | **Length** |
| --- | --- | --- | --- |
| 1Fnew | forward | AATGATACGGCGACCACCGAGATCTACACAAGCAGGATCGTCGGCAGC | 48 |
| 2Fnew | forward | AATGATACGGCGACCACCGAGATCTACACTTCGTCCTTCGTCGGCAGC | 48 |
| 3Fnew | forward | AATGATACGGCGACCACCGAGATCTACACCCTACTTCTCGTCGGCAGC | 48 |
| 4Fnew | forward | AATGATACGGCGACCACCGAGATCTACACGGATGAAGTCGTCGGCAGC | 48 |
| 5Fnew | forward | AATGATACGGCGACCACCGAGATCTACACAACAGCAATCGTCGGCAGC | 48 |
| 6Fnew | forward | AATGATACGGCGACCACCGAGATCTACACTTGTCGTTTCGTCGGCAGC | 48 |
| 7Fnew | forward | AATGATACGGCGACCACCGAGATCTACACGTTGATGGTCGTCGGCAGC | 48 |
| 8Fnew | forward | AATGATACGGCGACCACCGAGATCTACACAGAGTGTTTCGTCGGCAGC | 48 |
| 9Fnew | forward | AATGATACGGCGACCACCGAGATCTACACTCTCACAATCGTCGGCAGC | 48 |
| 10Fnew | forward | AATGATACGGCGACCACCGAGATCTACACAAGACACCTCGTCGGCAGC | 48 |
| 11Fnew | forward | AATGATACGGCGACCACCGAGATCTACACCAGGTAGTTCGTCGGCAGC | 48 |
| 12Fnew | forward | AATGATACGGCGACCACCGAGATCTACACACTTGCTGTCGTCGGCAGC | 48 |
| 13Rev_new | Reverse | CAAGCAGAAGACGGCATACGAGATGTCGTTCAGTCTCGTGGGCT | 44 |
| 14Rev_new | Reverse | CAAGCAGAAGACGGCATACGAGATATCTGGACGTCTCGTGGGCT | 44 |
| 15Rev_new | Reverse | CAAGCAGAAGACGGCATACGAGATGTATAGCGGTCTCGTGGGCT | 44 |
| 16Rev_new | Reverse | CAAGCAGAAGACGGCATACGAGATCATATCGCGTCTCGTGGGCT | 44 |
| 17Rev_new | Reverse | CAAGCAGAAGACGGCATACGAGATTATGCGACGTCTCGTGGGCT | 44 |
| 18Rev_new | Reverse | CAAGCAGAAGACGGCATACGAGATATGATACCGTCTCGTGGGCT | 44 |
| 19Rev_new | Reverse | CAAGCAGAAGACGGCATACGAGATTACTATGGGTCTCGTGGGCT | 44 |
| 20Rev_new | Reverse | CAAGCAGAAGACGGCATACGAGATAGTGGTAGGTCTCGTGGGCT | 44 |

6C: List of barcodes of table 6B

| **Plate row** | **Reverse primer** | **MiSeq I7 index used in sample sheet** | **Notes** |
| --- | --- | --- | --- |
| A | 13Rev_new | **TGAACGAC** | Designed in-house for this study |
| B | 14Rev_new | **GTCCAGAT** | Designed in-house for this study |
| C | 15Rev_new | **CGCTATAC** | Designed in-house for this study |
| D | 16Rev_new | **GCGATATG** | Designed in-house for this study |
| E | 17Rev_new | **GTCGCATA** | Designed in-house for this study |
| F | 18Rev_new | **GGTATCAT** | Designed in-house for this study |
| G | 19Rev_new | **CCATAGTA** | Designed in-house for this study |
| H | 20Rev_new | **CTACCACT** | Designed in-house for this study |
| **Plate column** | **Forward primer** | **MiSeq I5/index2 used in sample sheet** |  |
| 1 | 1Fnew | **AAGCAGGA** | Designed in-house for this study |
| 2 | 2Fnew | **TTCGTCCT** | Designed in-house for this study |
| 3 | 3Fnew | **CCTACTTC** | Designed in-house for this study |
| 4 | 4Fnew | **GGATGAAG** | Designed in-house for this study |
| 5 | 5Fnew | **AACAGCAA** | Designed in-house for this study |
| 6 | 6Fnew | **TTGTCGTT** | Designed in-house for this study |
| 7 | 7Fnew | **GTTGATGG** | Designed in-house for this study |
| 8 | 8Fnew | **AGAGTGTT** | Designed in-house for this study |
| 9 | 9Fnew | **TCTCACAA** | Designed in-house for this study |
| 10 | 10Fnew | **AAGACACC** | Designed in-house for this study |
| 11 | 11Fnew | **CAGGTAGT** | Designed in-house for this study |
| 12 | 12Fnew | **ACTTGCTG** | Designed in-house for this study |

### 7. Primer stock preparation

Prepare 100 µM stock solutions according to supplier yield. Prepare 5 µM working stocks for indexed primers by mixing 20 µL of 100 µM primer with 380 µL nuclease-free water. Label tubes clearly and store at -20 ^o^C.

Prepare the DR primer mix according to the validated notebook formulation: forward (F) DR components at 2 µM each plus 0.5 µM mDRsp15; reverse (R) DR components at 2 µM each. The real-run sheet denotes the combined DR primer mix as approximately F 8.5 µM and R 8 µM.

### 8. 96-well plate and index layout

The 96-well layout uses reverse indexed primers 13Rev_new–20Rev_new by rows A–H and forward indexed primers 1Fnew–12Fnew by columns 1–12. The MiSeq sample sheet must use the same row/column index combinations. From the 5 µM dilution use multichannel pipette and add according to the scheme 2 µl of each primer to the PCR plate (total of 4 µl for each well). Don’t forget to change the tips to prevent barcode mixing when it is necessary (for example when adding A-H barcodes to columns 1-12).

**Example of 96 wells plate**

| **Row** | **1** | **2** | **3** | **4** | **5** | **6** | **7** | **8** | **9** | **10** | **11** | **12** |
| --- | --- | --- | --- | --- | --- | --- | --- | --- | --- | --- | --- | --- |
| A | 11-22 | 143-22 | 283-22 | 333-22 | 360-22 | 394-22 | 415-22 | 452-22 | 365-22 | 86-22 | 497-22 | 525-22 |
| B | 24-22 | 264-22 | 285-22 | 334-22 | 363-22 | 396-22 | 416-22 | 461-22 | 375-22 | 94-22 | 498-22 | 529-22 |
| C | 37-22 | 271-22 | 288-22 | 335-22 | 367-22 | 400-22 | 417-22 | 462-22 | 243-22 | 114-22 | 504-22 | 532-22 |
| D | 41-22 | 274-22 | 289-22 | 336-22 | 372-22 | 401-22 | 420-22 | 464-22 | 211-22 | 73-22 | 510-22 | 547-22 |
| E | 43-22 | 276-22 | 311-22 | 344-22 | 373-22 | 404-22 | 422-22 | 470-22 | 322-22 | 163-22 | 515-22 | 554-22 |
| F | 44-22 | 277-22 | 316-22 | 345-22 | 387-22 | 408-22 | 428-22 | 471-22 | 8-22 | 130-22 | 516-22 | 556-22 |
| G | 48-22 | 278-22 | 317-22 | 355-22 | 388-22 | 413-22 | 429-22 | 477-22 | 76-22 | 332-22 | 523-22 | PTC |
| H | 70-22 | 279-22 | 320-22 | 359-22 | 389-22 | 414-22 | 430-22 | 364-22 | 78-22 | 495-22 | 524-22 | NTC |

### 9. Reaction setup

Each final PCR reaction is 20 µL. The indexed primers are pre-loaded into the plate; therefore the common master mix excludes the indexed primers and DNA template.

| **Component** | **Volume per reaction (µL)** | **Comment** |
| --- | --- | --- |
| Indexed forward primer, 5 µM | 2.0 | Pre-dispensed by column; final 0.5 µM |
| Indexed reverse primer, 5 µM | 2.0 | Pre-dispensed by row; final 0.5 µM |
| Nuclease-free water | 6.9 | Common master mix |
| 10× KOD buffer | 2.0 | Common master mix |
| dNTP mix, 2 mM | 2.0 | Common master mix |
| MgCl₂, 25 mM | 1.2 | Common master mix |
| DR primer mix | 0.5 | Common master mix; final DR Forward approx. 0.2125 µM; final Reverse 0.2 µM |
| ResoLight 480 dye, 20× | 1.0 | Common master mix |
| KOD Hot Start polymerase | 0.4 | Common master mix |
| DNA template | 2.0 | Add last |
| **Total** | **20.0** |  |

Procedure:

1. Use multichannel pipette and place 2 µL of each reverse indexed primer into all wells of the corresponding row.
2. Use multichannel pipette and place 2 µL of each forward indexed primer into all wells of the corresponding column.
3. Prepare common master mix for the number of reactions plus at least 10% excess.
4. Dispense 14 µL common master mix into each well already containing 4 µL of indexed primers.
5. Add 2 µL DNA template, positive (PTC) and nuclease-free water (NTC) control to the control wells.
6. Seal carefully with high-quality film or optical caps, centrifuge briefly, and transfer to the PCR instrument.

### 10. Thermal cycling program

| **Stage** | **Step** | **Temperature** | **Time** | **Cycles / notes** |
| --- | --- | --- | --- | --- |
| Initial denaturation | 1 | 95°C | 2 min |  |
| Denaturation | 2 | 95°C | 20 s |  |
| Annealing 1 | 3 | 64°C | 10 s |  |
| Extension & read | 4 | 70°C | 10 s | Toal 25 cycles, steps 2–4 |
| Denaturation | 5 | 95°C | 20 s |  |
| Annealing 2 | 6 | 50°C | 10 s |  |
| Extension/read | 7 | 70°C | 10 s | Total 12 cycles, steps 5–7 |
| Melt curve | 8 | 65–95°C | 0.5°C increments, 5 s each | Plate read of melting curve |
| Hold | 9 | 10°C | forever |  |

### 11. Pooling, cleanup and library QC

- After PCR, collect 10 µL from each well into a single pool in a trough using a multichannel pipette.
- Clean the pooled library using the ProNex kit at a 2× ratio or the laboratory-validated equivalent cleanup protocol.
- Quantify dsDNA concentration and perform smear/fragment analysis. In the real run, smear analysis was evaluated in the range up to 1500 bp.
- Record concentration, average size and calculated molarity in the library dilution worksheet.
- Reject or repeat libraries with abnormal background, poor concentration, unexpected size distribution or evidence of negative-control amplification.

### 12. Library dilution and MiSeq loading

| **Step** | **Instruction** | **Comment** |
| --- | --- | --- |
| Molarity formula | nM = [ng/µL] × 1,000,000 / (649 × average size bp) | Matches uploaded dilution worksheet. |
| Normalize to 4 nM | Volume of library for 10 µL = 4 / nM × 10 | Add RSB to 10 µL. |
| Denature | 5 µL pooled library + 5 µL 0.2 N NaOH; vortex and incubate 5 min | From uploaded dilution worksheet. |
| Dilute | Mix 990 µL Illumina HT1 buffer with 10 µL denatured library to 20 pM; dilute to 8-10 pM in a total volume of 600 µL. |  |
| PhiX | Add 6 µL of 20 pM PhiX (approx. 1%) | Load 606 µL of the library to MISeq reagent cartridge. |

### 13. MiSeq sample sheet

The standalone CSV file supplied with this package is a corrected example based on the real run. The template protects hyphenated sample IDs as text to prevent Excel date conversion, such as 11-22 becoming Nov-22.

| **Sample sheet field** | **Value used in example** |
| --- | --- |
| Module | GenerateFASTQ - 2.0.1 |
| Workflow | GenerateFASTQ |
| Library Prep Kit | Nextera DNA |
| Chemistry | Amplicon |
| Reads | 151 / 151 |
| reversecomplement | 0 |
| adapter | CTGTCTCTTATACACATCT |

### 14. Run acceptance and troubleshooting

| **Issue** | **Indicator** | **Action** |
| --- | --- | --- |
| Negative controls | Very low file size and spacer counts; no convincing spoligotype pattern | Investigate contamination if control has strong reads or recognizable profile. |
| Melting curve anomalies | Usually all MTC curves are cluster together. If not, consider repeating PCR or NTM contamination. | Repeat std-PCR |
| Low-output sample | R1+R2 approximately <1000 KB or mean spacer counts near background | Reject/repeat. Real run comments flagged 78-22 and 322-22. |
| Borderline output | R1 <1500 KB but R1+R2 >1000–2000 KB and coherent spacer pattern | May be informative; repeat if the result is important. |
| Strong MTBC test but absent/weak spacers | Possible rare DR-locus deletion or technical failure | Confirm with WGS/RD/SNP markers or another validated method. |
| Sample ID converted to a date | Hyphenated ID appears as month-year in sample sheet | Format all sample-ID cells as Text before entering IDs; use the Excel template. |

### 15. Records to retain

- Completed reaction-planning Excel file
- Primer lot and dilution records
- PCR run file and melt/amplicon review output
- Library QC report and dilution worksheet
- Final MiSeq sample sheet CSV
- MiSeq run folder and FASTQ files
- HiSpoT analysis output workbook
- Deviation/repeat notes

### 16. Appendix: full primer table

The Excel template also contains the primer order list and barcode map. Use the Excel file as the controlled operational document.
